# Evolution of a plant gene cluster in Solanaceae and emergence of metabolic diversity

**DOI:** 10.1101/2020.03.04.977231

**Authors:** Pengxiang Fan, Peipei Wang, Yann-Ru Lou, Bryan J. Leong, Bethany M. Moore, Craig A. Schenck, Rachel Combs, Pengfei Cao, Federica Brandizzi, Shin-Han Shiu, Robert L. Last

**Author notes:** Center for Applied Plant Sciences, The Ohio State University, Columbus, OH 43210.

## Abstract

Plants produce phylogenetically and spatially restricted, as well as structurally diverse specialized metabolites via multistep metabolic pathways. Hallmarks of specialized metabolic evolution include enzymatic promiscuity, recruitment of primary metabolic enzymes and genomic clustering of pathway genes. Solanaceae plant glandular trichomes produce defensive acylsugars, with aliphatic sidechains that vary in length across the family. We describe a tomato gene cluster on chromosome 7 involved in medium chain acylsugar accumulation due to trichome specific acyl-CoA synthetase and enoyl-CoA hydratase genes. This cluster co-localizes with a tomato steroidal alkaloid gene cluster forming a ‘supercluster’, and is syntenic to a chromosome 12 region containing another acylsugar pathway gene. We reconstructed the evolutionary events leading to emergence of this gene cluster and found that its phylogenetic distribution correlates with medium chain acylsugar accumulation across the Solanaceae. This work reveals dynamics behind emergence of novel enzymes from primary metabolism, gene cluster evolution and cell-type specific metabolite diversity.

## Introduction

Despite the enormous structural diversity of plant specialized metabolites, they are derived from a relatively small number of primary metabolites, such as sugars, amino acids, nucleotides, and fatty acids (*1*). These lineage-, tissue- or cell-type specific specialized metabolites mediate environmental interactions, such as herbivore and pathogen deterrence or pollinator and symbiont attraction (*2, 3*). Specialized metabolism evolution is primarily driven by gene duplication (*4, 5*), and relaxed selection of the resulting gene pairs allows modification of cell- and tissue-specific gene expression and changes in enzymatic activity. This results in expanded substrate recognition and/or diversified product formation (*6, 7*). The neofunctionalized enzymes can prime the origin and diversification of specialized metabolic pathways (*8–10*).

There are many examples of mechanisms that lead to novel enzymatic activities in specialized cell or tissue types, however, the principles that govern assembly of multi-enzyme specialized metabolic pathways are less well established. One appealing hypothesis involves the stepwise recruitment of pathway enzymes (*11*). Clustering of non-homologous specialized metabolic enzyme genes, which participate in the same biosynthetic pathway, is an unusual feature of plant genomes (*12, 13*). This phenomenon is seen for pathways leading to products of diverse structures and biosynthetic origins including terpenes (*14, 15*), cyclic hydroxamic acids (*16*), alkaloids (*17, 18*), polyketides (*19*), cyanogenic glucosides (*20*), and modified fatty acids (*21*). However, while these clusters encode multiple non-homologous enzymes of a biosynthetic pathway, evolution of their assembly is not well understood.

Acylsugars are a group of insecticidal (*22*) and anti-inflammatory (*23*) chemicals mainly observed in glandular trichomes of Solanaceae species (*24, 25*). These specialized metabolites are sugar aliphatic esters with three levels of structural diversity across the Solanaceae family: acyl chain length, acylation position, and sugar core (*24*). The primary metabolites sucrose and aliphatic acyl-CoAs are the biosynthetic precursors of acylsucroses in plants as evolutionarily divergent as the cultivated tomato *Solanum lycopersicum* (*26*) (Fig. 1), *Petunia axillaris* (*27*) and *Salpiglossis sinuata* (*28*). The core tomato acylsucrose biosynthetic pathway involves four BAHD [**B**EAT, **A**HCT, **H**CBT, **D**AT(*29*)] family acylsucrose acyltransferases (*Sl-ASAT1* through *Sl-ASAT4*), which are specifically expressed in the type I/IV trichome tip cells (*26, 30, 31*). These enzymes catalyze consecutive reactions utilizing sucrose and acyl-CoA substrates to produce the full set of cultivated tomato acylsucroses *in vitro* (*26*).

**Fig. 1.**
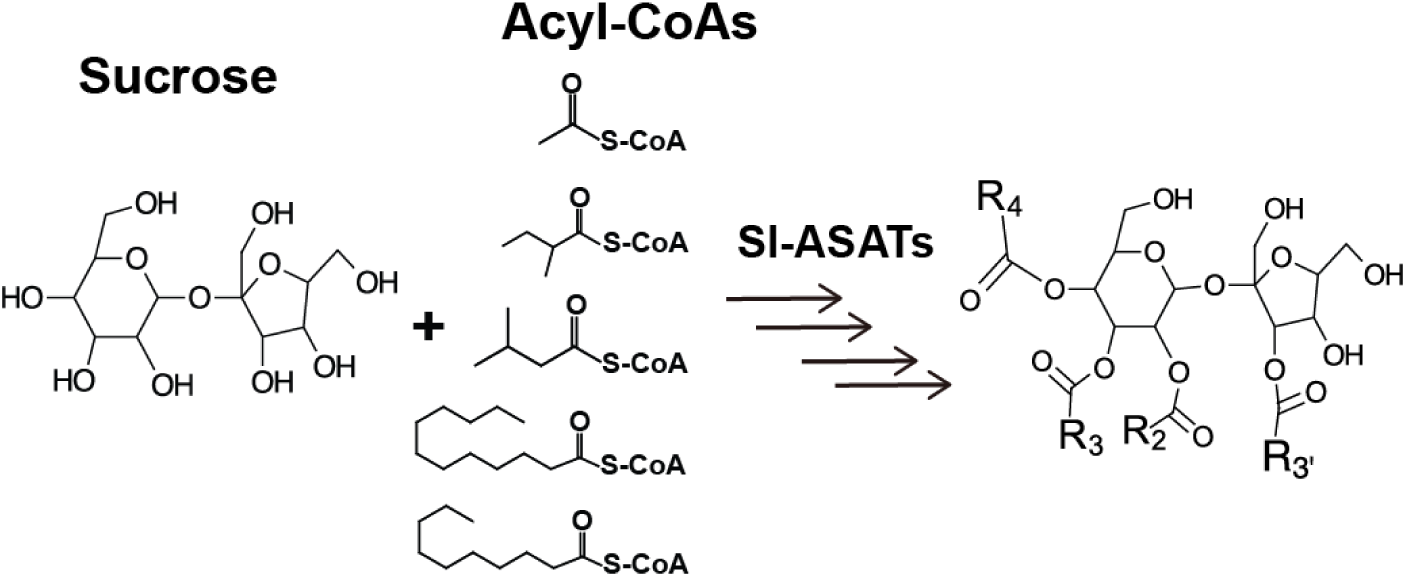
Primary metabolites are biosynthetic precursors of tomato trichome acylsugars. In cultivated tomatoes, the trichome acylsucroses are synthesized by four Sl-ASATs using the primary metabolites – sucrose and different types of acyl-CoAs – as substrates.

Co-option of primary metabolic enzymes contributed to the evolution of acylsugar biosynthesis and led to interspecific structural diversification across *Solanum* tomato clade. One example is an invertase-like enzyme originating from carbohydrate metabolism that generates acylglucoses in the wild tomato *S. pennellii* through cleavage of the acylsucrose glycosidic bond (*32*). In another case, allelic variation of a truncated isopropylmalate synthase-like enzyme (IPMS3) – from branched chain amino acid metabolism – leads to acylsugar iC4/iC5 (2-methylpropanoic/3-methylbutanoic acid) acyl chain diversity in *S. pennellii* and *S. lycopersicum* (*33*). Acylsugar structural diversity is even more striking across the family. Previous studies revealed variation in acyl chain length (*28, 34, 35*): *Nicotiana*, *Petunia* and *Salpiglossis* species were reported to accumulate acylsugars containing only short acyl chains (carbon number, C ≤ 8). In contrast, some species in *Solanum* and other closely related genera produce acylsugars with medium acyl chains (C ≥ 10). These results are consistent with the hypothesis that the capability to produce medium chain acylsugars varies across the Solanaceae family.

In this study, we identify a metabolic gene cluster on tomato chromosome 7 containing two non-homologous genes – acylsugar acyl-CoA synthetase (*AACS*) and acylsugar enoyl-CoA hydratase (*AECH*) – affecting medium chain acylsugar biosynthesis. Genetic and biochemical results show that the trichome enriched *AACS* and *AECH* are involved in generating medium chain acyl-CoAs, which are donor substrates for acylsugar biosynthesis. Genomic analysis revealed a syntenic region of the gene cluster on chromosome 12, where the acylsucrose biosynthetic *Sl-ASAT1* is located (*26*). Phylogenetic analysis of the syntenic regions in Solanaceae and beyond led to evolutionary reconstruction of the origin of the acylsugar gene cluster. We infer that sequential gene insertion facilitated emergence of this gene cluster, while gene duplication and pseudogenization rearranged and fragmented it. These results provide insights into how co-option of primary metabolic enzymes, emergence of cell-type specific gene expression, and the formation of metabolic gene clusters play a role in specialized metabolism evolution.

## Results

### Identification of a metabolic gene cluster that affects tomato trichome medium chain acylsugar biosynthesis

*S. pennellii* natural accessions (*36*), as well as the *S. lycopersicum* M82 × *S. pennellii* LA0716 chromosomal substitution introgression lines (ILs) (*37*), offer convenient resources to investigate interspecific genetic variation that affects acylsugar metabolic diversity (*36, 38*). In a rescreen of ILs for *S. pennellii* genetic regions that alter trichome acylsugar profiles (*38*), IL7-4 was found to accumulate increased C10 medium chain containing acylsugars compared with M82 (Fig. 2, A and B). The genetic locus that contributes to the acylsugar phenotype was narrowed down to a 685 kb region through screening selected backcross inbred lines (BILs) (*39*) that have recombination breakpoints on chromosome 7 (Fig. 2C). Because tomato acylsucrose biosynthesis occurs in trichomes, candidate genes in this region were filtered based on their trichome-specific expression patterns. This analysis identified a locus containing multiple tandem duplicated genes of three types – acyl-CoA synthetases (ACS), enoyl-CoA hydratases (ECH), and BAHD family acyltransferases. Three genes with expression enriched in tomato trichomes were selected for further analysis (Fig. 2D).

**Fig. 2.**
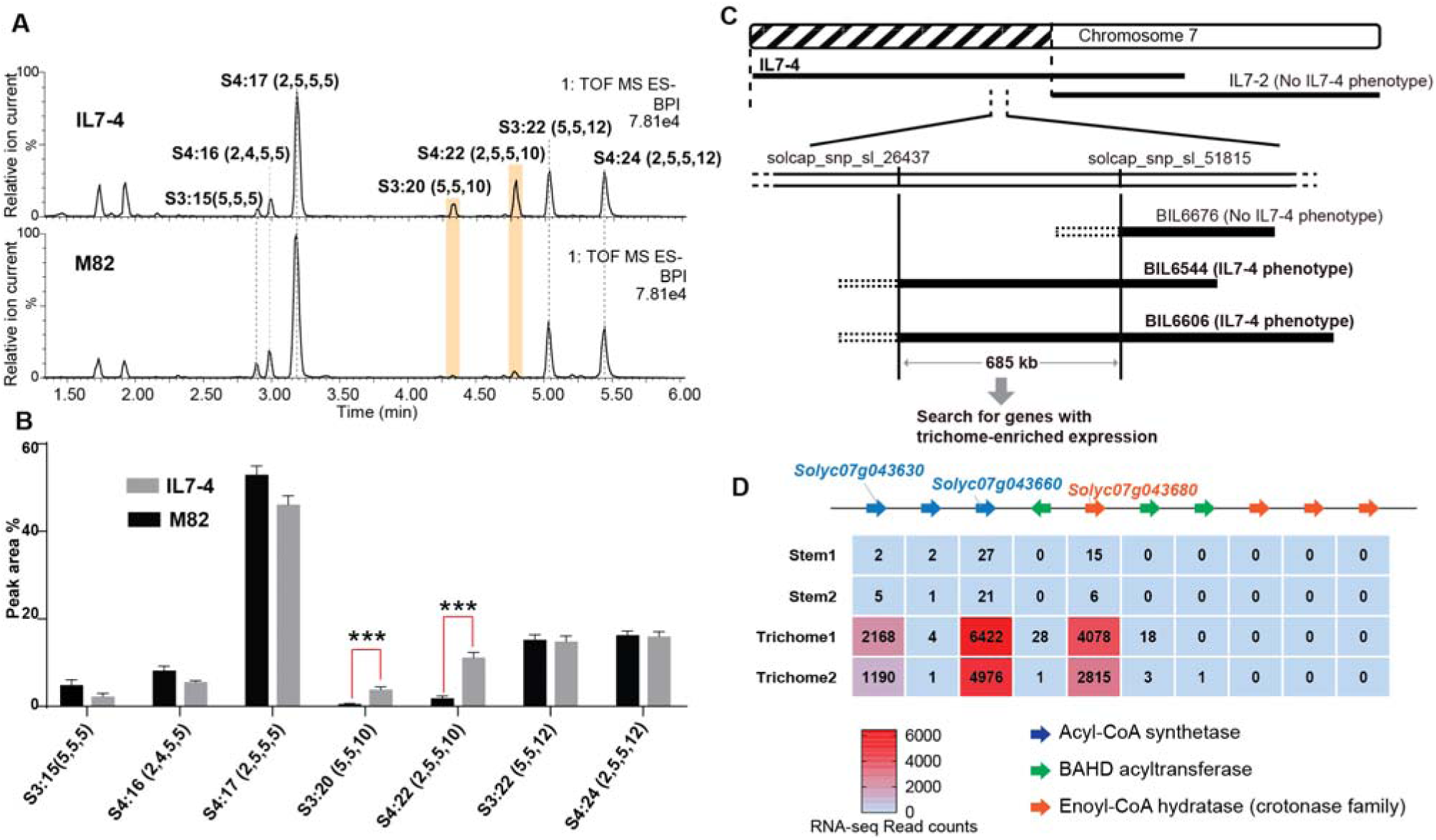
Mapping of a genetic locus related to acylsugar variations in tomato interspecific introgression lines. (**A**) Electrospray ionization negative (ESI^-^) mode, base-peak intensity (BPI) LC/MS chromatogram of trichome metabolites from cultivated tomato *S. lycopersicum* M82 and introgression line IL7-4. The orange bars highlight two acylsugars that have higher abundance in IL7-4 than in M82. For the acylsucrose nomenclature, “S” refers to a sucrose backbone, “3:22” means three acyl chains with twenty-two carbons in total. The length of each acyl chain is shown in the parentheses. (**B**) Peak area percentage of seven major trichome acylsugars in M82 and IL7-4. The sum of the peak area percentage of each acylsugar is equal to 100% in each sample. ***p < 0.001, Welch two-sample *t* test, n=3. (**C**) Mapping the genetic locus contributing to the IL7-4 acylsugar phenotype using selected backcross inbred lines (BILs) that have recombination break points within the introgression region of IL7-4. (**D**) Narrowing down candidate genes in the locus using trichome/stem RNA-seq datasets generated from previous study (*33*). A region with duplicated genes of three types – acyl-CoA synthetase (ACS), BAHD acyltransferase, and enoyl-CoA hydratase (ECH) – is shown. The red-blue color gradient provides a visual marker to rank the expression levels represented by read counts.

The three candidate genes were tested for involvement in tomato acylsugar biosynthesis by making loss of function mutations using the CRISPR-Cas9 gene editing system. Two guide RNAs (gRNAs) were designed to target one or two exons of each gene to assist site-specific DNA cleavage by hCas (*40*) (fig. S1, A-C). In the self-crossed T1 progeny of stably transformed M82 plants, at least two homozygous mutants were obtained in *Solyc07g043630*, *Solyc07g043660*, and *Solyc07g043680* (fig. S1, A-C), and these were analyzed for leaf trichome acylsugar changes. Altered acylsugar profiles were observed in the ACS-annotated *Solyc07g043630* or ECH-annotated *Solyc07g043680* mutants (Fig. 3, A and B), but not in the ACS-annotated *Solyc07g043660* mutant (fig. S1D). Despite carrying mutations in distinctly annotated genes (ACS or ECH), the two mutants exhibited the same phenotype – no detectable medium acyl chain (C10 or C12) containing acylsugars (Fig. 3, A and B). We renamed *Solyc07g043630* as *<u>a</u>cylsugar <u>a</u>cyl-<u>C</u>oA <u>s</u>ynthetase 1* (*Sl-AACS1*) and *Solyc07g043680* as *<u>a</u>cylsugar <u>e</u>noyl-<u>C</u>oA <u>h</u>ydratase 1* (*Sl-AECH1*) based on this analysis.

**Fig. 3.**
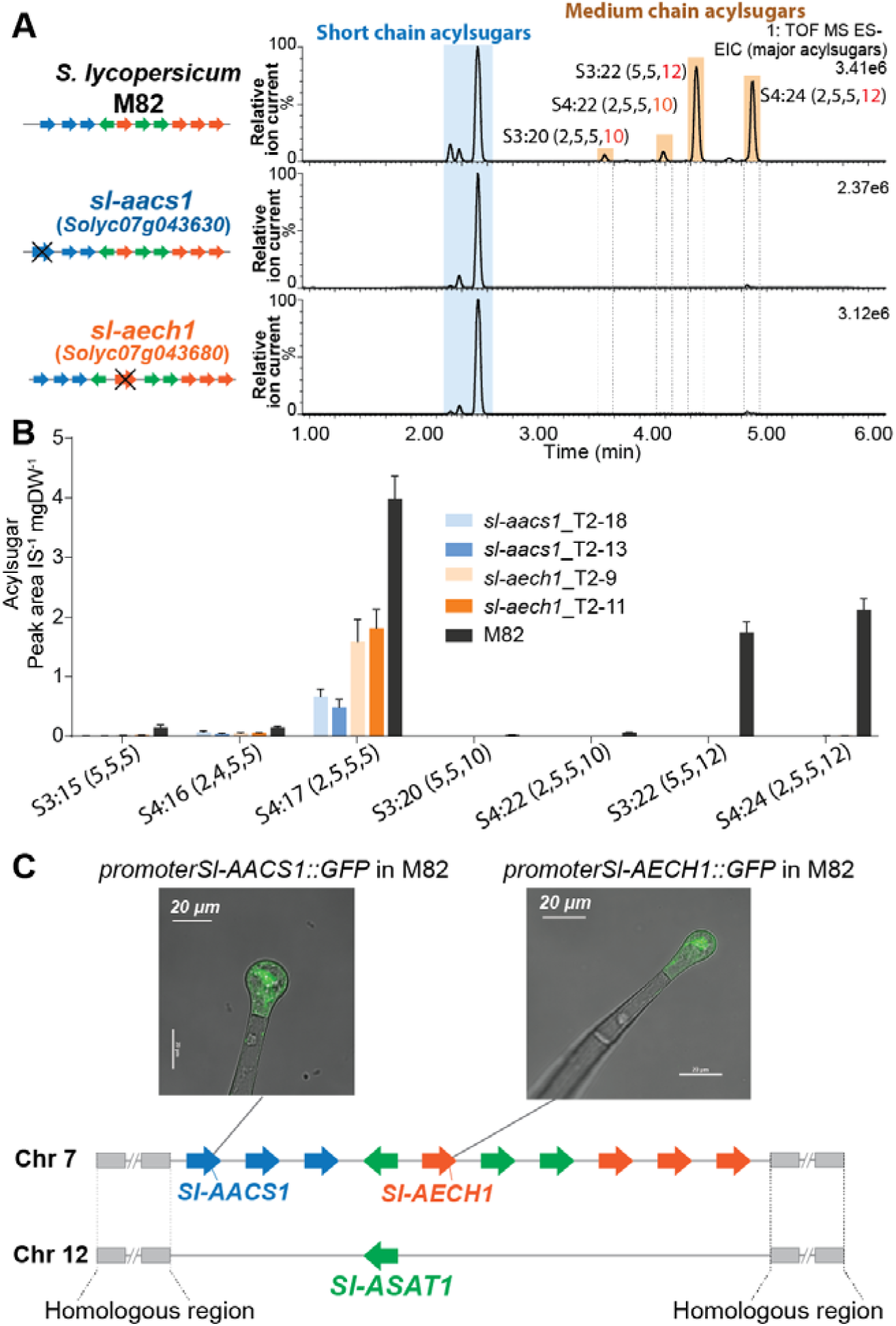
CRISPR/Cas9-mediated gene knockout of tomato *Sl-AACS1* or *Sl-AECH1* eliminates detectable medium chain containing acylsugars. (**A**) Combined LC/MS extracted ion chromatograms of trichome metabolites from CRISPR mutants *sl-aacs1* and *sl-aech1.* The medium chain acylsugars that are not detected in the two mutants are denoted by pairs of vertical dotted lines. (**B**) Quantification of seven major trichome acylsugars in *sl-aacs1* and *sl-aech1* mutants. Two independent T2 generation transgenic lines for each mutant were used for analysis. The peak area/internal standard (IS) normalized by leaf dry weight (DW) is shown from six biological replicates ± SE. (**C**) Confocal fluorescence images showing that GFP fluorescence driven by *Sl-AACS1* or *Sl-AECH1* is located in the tip cells of type I/IV trichomes. Their tissue specific expressions are similar to *Sl-ASAT1* (*26*), which locates in a chromosome 12 region that is syntenic to the locus containing *Sl-AACS1* and *Sl-AECH1*.

Further genomic analysis revealed that *Sl-AACS1* and *Sl-AECH1* belong to a syntenic region shared with a locus on chromosome 12, where *Sl-ASAT1* is located (Fig. 3C and fig. S2)*. Sl-ASAT1* is specifically expressed in trichome tip cells and encodes the enzyme catalyzing the first step of tomato acylsucrose biosynthesis (*26*). This led us to test the cell-type expression pattern of *Sl-AACS1* and *Sl-AECH1.* Like *Sl-ASAT1*, the promoters of both genes drove GFP expression in the trichome tip cells of stably transformed M82 plants (Fig. 3C). This supports our hypothesis that *Sl-AACS1* and *Sl-AECH1* are involved in tomato trichome acylsugar biosynthesis. Taken together, we identified a metabolic gene cluster involved in medium chain acylsugar biosynthesis, which is composed of two cell-type specific genes.

### *In vitro* analysis of Sl-AACS1 and Sl-AECH1 implicates their roles in medium chain acyl-CoA metabolism

ACS and ECH are established to function in multiple cell compartments for the metabolism of acyl-CoA(*41*), the acyl donor substrates for ASAT enzymes. We sought to understand the organelle targeting of Sl-AACS1 and Sl-AECH1, to advance our knowledge of acylsugar machinery at the subcellular level. We constructed expression cassettes of Sl-AACS1, Sl-AECH1 and Solyc07g043660 with C-terminal cyan fluorescent protein (CFP), hypothesizing that the targeting peptides reside at the N-terminus of precursor proteins. When co-expressed in tobacco leaf epidermal cells, three CFP-tagged recombinant proteins co-localized with the mitochondrial marker MT-RFP (*42*) (Fig. 4A and fig. S3A). To rule out the possibility of peroxisomal localization, we fused Sl-AACS1, Sl-AECH1, or Solyc07g043660 with N-terminus fused yellow fluorescent protein (YFP), considering that potential peroxisomal targeting peptides are usually located on the C-terminus (*43*). The expressed YFP-recombinant proteins were not co-localized with the peroxisomal marker RFP-PTS (*42*) (fig. S3B). Instead, they appeared distributed in the cytosol (fig. S3B), presumably because the N-terminal YFP blocked the mitochondria targeting signal. Taken together, protein expression and co-localization analyses suggest that Sl-AACS1, Sl-AECH1, and Solyc07g043660 encode enzymes targeted to mitochondria.

**Fig. 4.**
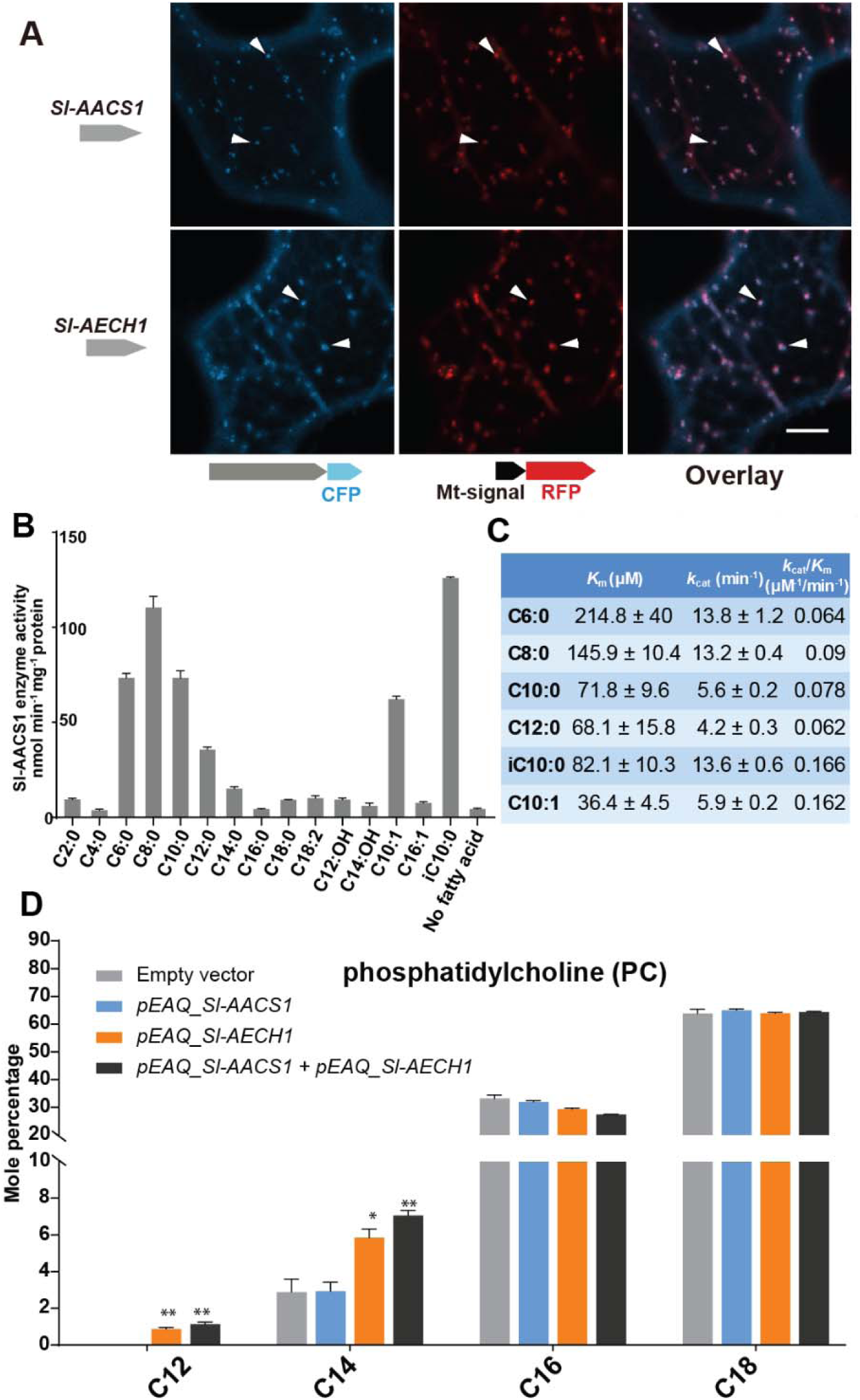
Functional analysis of Sl-AACS1 and Sl-AECH1 in *N. benthamiana* and recombinant Sl-AACS1 enzyme analysis. (**A**) Confocal images of co-expression analysis in tobacco leaf epidermal cells using C-terminal CFP-tagged either Sl-AACS1 or Sl-AECH1 and the mitochondrial marker MT-RFP. Arrowheads point to mitochondria that are indicated by MT-RFP fluorescent signals. Scale bar equals 10 μm. (**B**) Aliphatic fatty acids of different chain lengths were used as the substrates to test Sl-AACS1 acyl-CoA synthase activity. Mean amount of acyl-CoAs generated (nmol min^-1^ mg^-1^ proteins) was used to represent enzyme activities. The results are from three measurements ± SE. (**C**) Enzyme activity of Sl-AACS1 for six fatty acid substrates. (**D**) Identification of membrane lipid phosphatidylcholine (PC), which contains medium acyl chains, following transient expression of *Sl-AECH1* in *N. benthamiana* leaves. The results from expressing *Sl-AACS1* and co-expressing both *Sl-AECH1* and *Sl-AACS1* are also shown. Mole percentage (Mol %) of the acyl chains from membrane lipids with carbon number 12, 14, 16, and 18 are shown for three biological replicates ± SE. *p < 0.05, **p < 0.01. Welch two-sample *t* test was performed comparing with the empty vector control. Acyl groups of the same chain lengths with saturated and unsaturated bonds were combined in the calculation.

Sl-AACS1 belongs to a group of enzymes that activate diverse carboxylic acid substrates to produce acyl-CoAs. We hypothesized that Sl-AACS1 uses medium chain fatty acids as substrates, because ablation of *Sl-AACS1* eliminated acylsugars with medium acyl chains. To characterize the *in vitro* activity of Sl-AACS1, we purified recombinant His-tagged proteins from *Escherichia coli*. Enzyme assays were performed by supplying fatty acid substrates with even carbon numbers from C2 through C18 (Fig. 4B). The results showed that Sl-AACS1 utilized fatty acid substrates with lengths ranging from C6 to C12, including those with a terminal branched carbon (iC10:0) or an unsaturated bond (*trans*-2-decenoic acid, C10:1) (Fig. 4, B and C). However, no activity was observed with the 3-hydroxylated C12 and C14 fatty acids as substrates (Fig. 4B). These results support our hypothesis that Sl-AACS1 produces medium chain acyl-CoAs, which are *in vivo* substrates for acylsugar biosynthesis.

To test whether *Sl-AACS1* and *Sl-AECH1* can produce medium chain acyl-CoAs *in planta,* we transiently expressed these genes in *Nicotiana benthamiana* leaves using *Agrobacterium*-mediated infiltration (*44*). It is challenging to directly measure plant acyl-CoAs, due to their low concentration and separate organellar pools. We used an alternative approach and characterized membrane lipids, which are produced from acyl-CoA intermediates. We took advantage of the observation that *N. benthamiana* membrane lipids do not accumulate detectable acyl chains of 12 carbons or shorter. *N. benthamiana* leaves were infiltrated with constructs containing *Sl-AACS1* or *Sl-AECH1* individually, or together (Fig. 4D). In contrast to the empty vector control, infiltration of *Sl-AECH1* led to detectable levels of C12 acyl chains in the leaf membrane lipid phosphatidylcholine (PC) (Fig. 4D). We also observed increased C14 acyl chains in PC, phosphatidylglycerol (PG), sulfoquinovosyl diacylglycerol (SQDG), and digalactosyldiacylglycerol (DGDG) in *Sl-AECH1* infiltrated plants (Fig. 4D and fig. S3C). These results suggest that *Sl-AECH1* participates in generation of medium chain acyl-CoAs *in planta*, which are channeled into lipid biosynthesis. No medium chain acylsugars were detected, presumably due to the lack of core acylsugar biosynthetic machinery in *N. benthamiana* mesophyll cells.

We asked whether the closest known homologs of *Sl-AECH1* from *Solanum* species can generate medium chain lipids when transiently expressed in *N. benthamiana*. Two SQDGs with C12 chains were monitored by LC/MS as peaks diagnostic of lipids containing medium chain fatty acids (fig. S4, A and B). The results showed that only the putative *Sl-AECH1* orthologs *Sopen07g023250* (*Sp-AECH1*) and *Sq_c37194* (*Sq-AECH1*) – from *S. pennellii* and *S. quitoense* respectively *–* generated medium chain lipids in the infiltrated leaves (fig. S4C). This confirms that not all ECHs can produce medium chain lipids and suggests that the function of *Sl-AECH1* evolved recently, presumably as a result of neofunctionalization after gene duplication (fig. S4C).

### *AACS1* and *AECH1* are evolutionarily conserved in the *Solanum*

Medium chain acylsugars were documented in *Solanum* species besides cultivated tomato, including *S. pennellii* (*32*), *S. nigrum* (*28*), as well as the more distantly related *S. quitoense* (*45*) and *S. lanceolatum* (*23*). We hypothesized that evolution of *AACS1* and *AECH1* contributed to medium chain acylsugar biosynthesis in *Solanum.* As a test, we analyzed the genomes of *Solanum* species other than cultivated tomato. Indeed, the acylsugar gene cluster containing ACS and ECH was found in both *S. pennellii* and *S. melongena* (eggplant), suggesting that the cluster assembly evolved before divergence of the tomato and eggplant lineage (Fig. 5A).

**Fig. 5.**
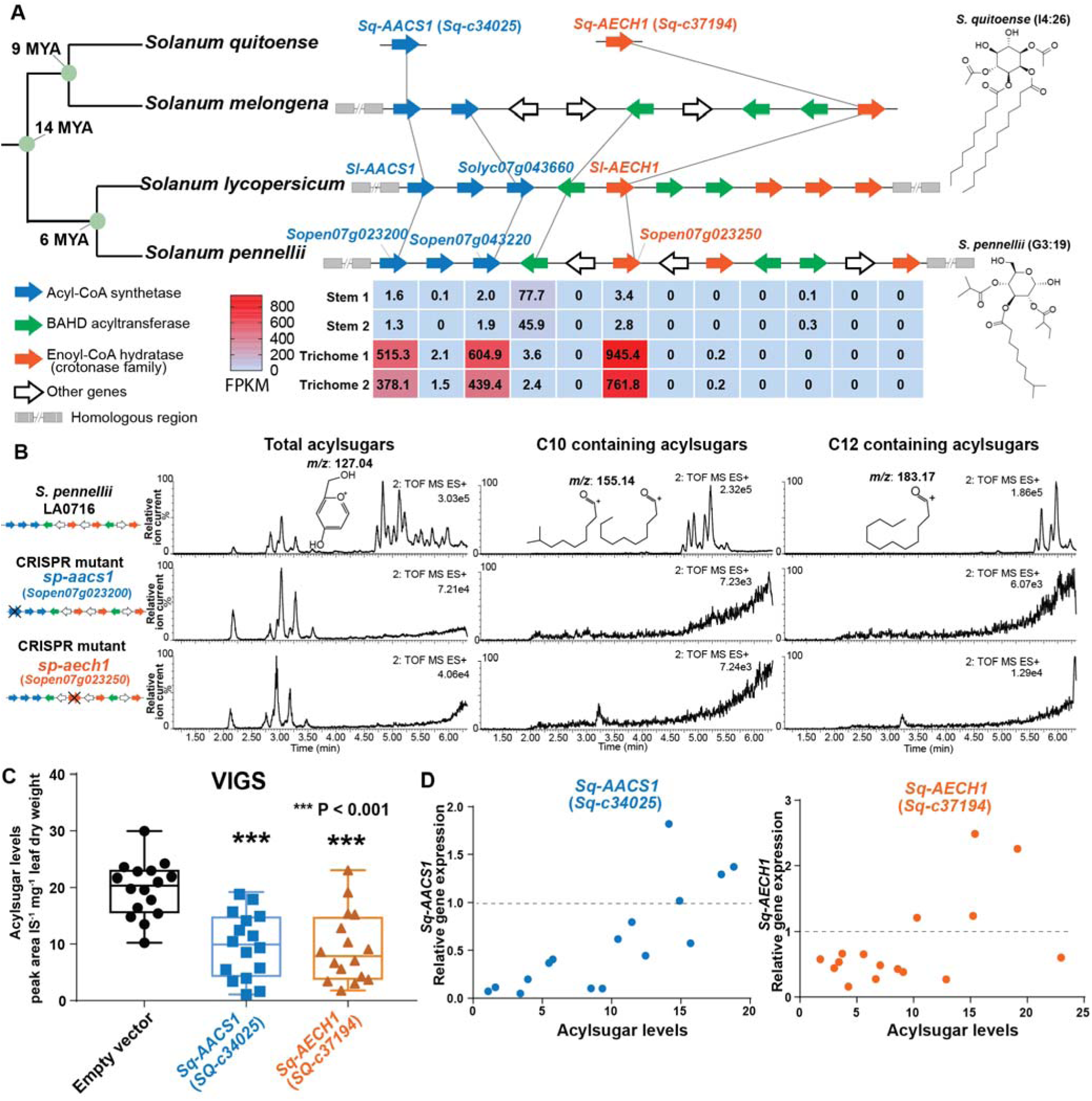
*AACS1* and *AECH1* are evolutionarily conserved in *Solanum* plants. (**A**) A conserved syntenic genomic region containing *AACS1* and *AECH1* was found in three selected *Solanum* species. Nodes representing estimated dates since the last common ancestors (*69*) shown on the left. The closest homologs of *AACS1* and *AECH1* in *Solanum quitoense* are shown without genomic context because the genes were identified from RNA-seq and genome sequences are not available. The lines connect genes representing putative orthologs across the four species. The trichome/stem RNA-seq data of two biological *S. pennellii* replicates are summarized (table S1) for genes in the syntenic region. The red-blue color gradient provides a visual marker to rank the expression levels in Fragments Per Kilobase of transcript per Million mapped reads (FPKM). Structures of representative medium chain acylsugars from *S. quitoense* (acylinositol, I4:26) (*45*) and *S. pennellii* (acylglucose, G3:19) (*32*) are on the right. (**B**) CRISPR/Cas9-mediated gene knockout of *Sp-AACS1* or *Sp-AECH1* in *S. pennellii* produce no detectable medium chain containing acylsugars. The ESI^+^ mode LC/MS extracted ion chromatograms of trichome metabolites are shown for each mutant. The *m/z* 127.01 (left panel) corresponds to the glucopyranose ring fragment that both acylsucroses and acylglucoses generate under high collision energy positive-ion mode. The *m/z* 155.14 (center panel) and 183.17 (right panel) correspond to the acylium ions from acylsugars with chain length of C10 and C12, respectively. (**C**) Silencing *Sq-AACS1* (*Sq-c34025*) or *Sq-AECH1* (*Sq-c37194*) in *S. quitoense* using VIGS leads to reduction of total acylsugars. The peak area/internal standard (IS) normalized by leaf dry weight was shown from fifteen biological replicates. ***p < 0.001, Welch two-sample *t* test. (**D**) Reduced gene expression of *Sq-AACS1* or *Sq-AECH1* correlates with decreased acylsugar levels in *S. quitoense*. The qRT-PCR gene expression data are plotted with acylsugar levels of the same leaf as described in Fig. S6E.

We applied gene expression and genetic approaches to test the *in vivo* functions of ACS and ECH in selected *Solanum* species. To explore the expression pattern of *S. pennellii* ACS and ECH cluster genes, we performed RNA-seq analysis on trichomes and shaved stems to identify acylsugar biosynthetic candidates (table S1). The expression pattern of *S. pennellii* cluster genes is strikingly similar to *S. lycopersicum*: one ECH and two ACS genes are highly enriched in trichomes, including the orthologs of *Sl-AACS1* and *Sl-AECH1*. *Sp-AACS1* function (*Sopen07g023200*) was first tested by asking whether it can reverse the cultivated tomato *sl-aacs1* mutant acylsugar phenotype. Indeed, *Sp-AACS1* restored C12 containing acylsugars in the stably transformed *sl-aacs1* plants (fig. S5). To directly test *Sp-AACS1* and *Sp-AECH1* function, we used CRISPR-Cas9 to make single mutants in *S. pennellii* LA0716. No medium chain acylsugars were detected in T0 generation mutants with edits for each gene (Fig. 5B and fig. S6, A and C). Similar to the ACS-annotated *Solyc07g043660* cultivated tomato mutant (fig. S1D), deletion of *S. pennellii* ortholog *Sopen07g023220* has no observed effects on *S. pennellii* trichome acylsugars (fig. S6D).

**Fig. 6.**
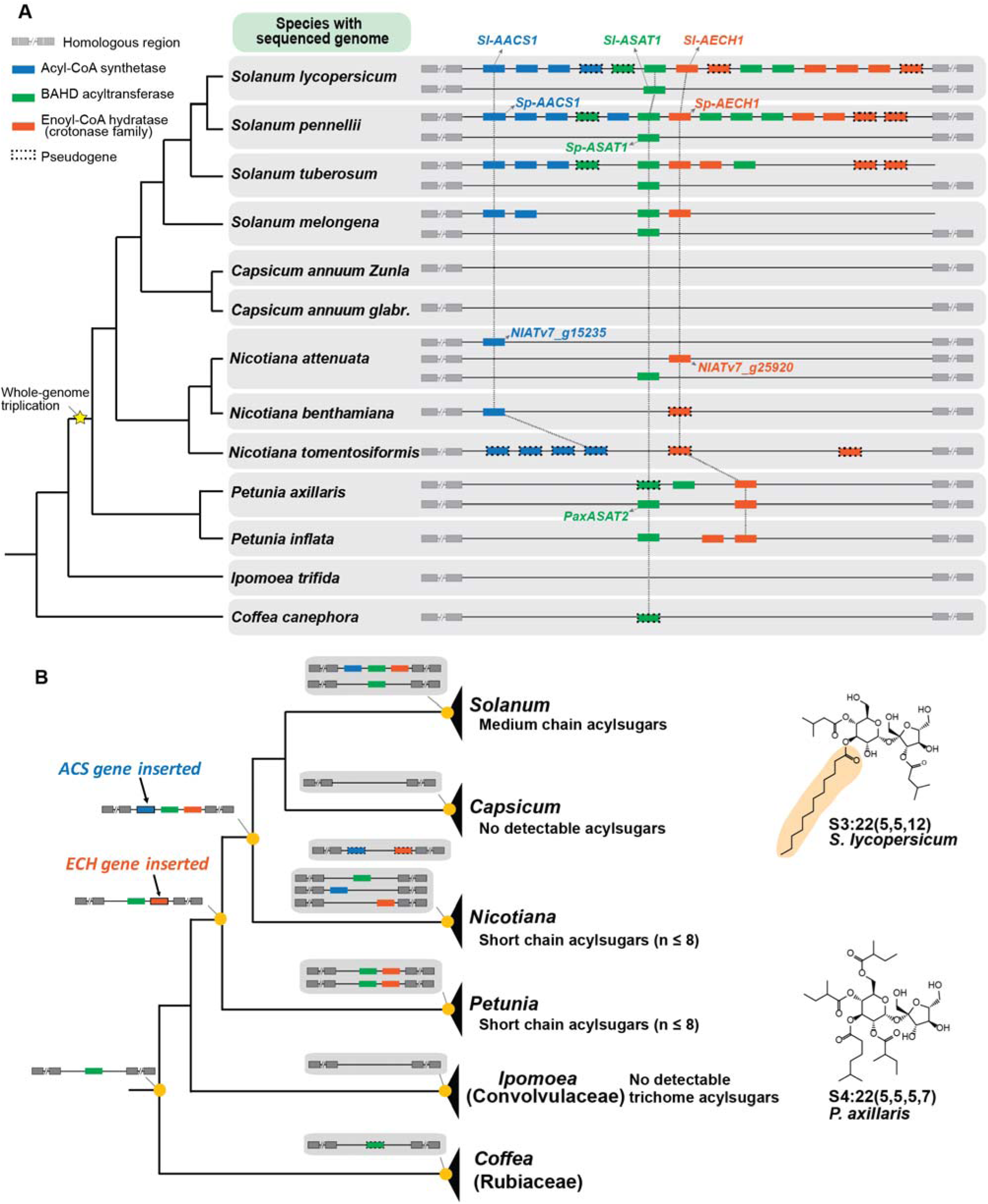
Evolution of the acylsugar gene cluster is associated with acylsugar acyl chain diversity across the Solanaceae family. (**A**) The acylsugar gene cluster syntenic regions of 11 Solanaceae species and two outgroup species *Ipomea trifida* (Convolvulaceae) and *Coffea canephora* (Rubiaceae). This is a simplified version adapted from fig. S7. Only genes from the three families – ACS (blue), BAHD acyltransferase (green), and ECH (orange) – are shown. The vertical lines connect putative orthologs. For information about the syntenic region size in each species refer to fig. S7. (**B**) The evolutionary history of the acylsugar gene cluster and its relation to the acylsugar phenotypic diversity. Structures of representative short and medium chain acylsugars were shown on the right.

The medium chain acylsugar producer *S. quitoense* (*45*) was used for *AACS1* and *AECH1* functional analysis because of its phylogenetic distance from the tomato clade, it is in the *Solanum* Leptostemonum clade (including eggplant), and the fact that it produces medium chain acylsugars. We found trichome-enriched putative orthologs of *AACS1* and *AECH1* in the transcriptome dataset of *S. quitoense* (*28*), and tested their *in vivo* function through virus-induced gene silencing (VIGS) (fig. S6E). Silencing either gene led to decreased total acylsugars (Fig. 5C and fig. S6F), which correlated with the degree of expression reduction in each sample (Fig. 5D). These results are consistent with the hypothesis that *Sq-AACS1* and *Sq-AECH1* are involved in medium chain acylsugar biosynthesis, because all acylsugars in *S. quitoense* carry two medium chains (*45*). The importance of *AACS1* and *AECH1* in medium chain acylsugar biosynthesis in distinct *Solanum* clades inspired us to explore the evolutionary origins of the gene cluster.

### Evolution of the gene cluster correlates with the distribution of medium chain acylsugars across Solanaceae

We sought to understand how the acylsugar gene cluster evolved and whether it correlates with the distribution of medium chain acylsugars across the Solanaceae family. Taking advantage of the available genome sequences of 13 species from Solanaceae and sister families, we analyzed the regions that are syntenic with the tomato acylsugar gene cluster (fig. S7). This synteny was found in all these plants, including the most distantly related species analyzed, *Coffea canephora* (Rubiaceae) (fig. S7). BAHD acyltransferase genes were the only cluster genes observed in the syntenic regions both inside and outside the Solanaceae, in contrast to ECH and ACS, which are restricted to the family (Fig. 6A and fig. S7). Within the syntenic regions of the species analyzed, ECH homologs, including pseudogenes, are present in all Solanaceae except for *Capsicum* species, while ACS is more phylogenetically restricted, being found only in *Nicotiana* and *Solanum* (Fig. 6A and fig. S7).

Based on the species phylogeny, we reconstructed the evolutionary history of the acylsugar gene cluster in Solanaceae according to the presence or absence of these three genes in the syntenic block (Fig. 6B). We propose two sequential gene insertion events that took place during the evolution of Solanaceae followed by gene duplications and gene losses leading to the cluster structures in extant plants (Fig. 6B). In this model, the BAHD acyltransferase was present in the most recent common ancestor of Solanaceae, Convolvulaceae, and Rubiaceae, and was likely lost in Ipomoea. An ancestral ECH gene insertion into the syntenic region prior to the divergence of *Petunia* and its sister clade was the next event. This was followed by the insertion of an ancestral ACS gene into the cluster in the most recent common ancestor of *Nicotiana, Capsicum*, and *Solanum* (Fig. 6B). The entire cluster was lost in *Capsicum* (Fig. 6B).

We propose that the emergence of both ACS and ECH genes in the synteny was a prerequisite for the rise of medium chain acylsugars in Solanaceae species (Fig. 6B). Consistent with the hypothesis, only short chain acylsugars were observed in *Petunia* (*35*) (fig. S8), which correlates with the absence of ACS homolog (Fig. 6B). In contrast, medium chain acylsugars were detected throughout *Solanum* (fig. S8), supported by the observation that both ACS and ECH homologs are present in extant *Solanum* species (Fig. 6B). Interestingly, although *Nicotiana* species collectively have both ACS and ECH genes (Fig. 6B), they do not produce medium chain acylsugars (fig. S8) due to differential gene losses in different species. For example, the ECH homologs are pseudogenes in *N. benthamiana* and *N. tomentosiformis* (Fig. 6A), and the ECH gene in *N. attenuata,* is not located within the same syntenic region with ACS gene (fig. S7), and has very low expression in all the tissues tested (*46*), including in trichome-rich tissues such as stems and pedicels (table S2). These results show that the presence of both functional ACS and ECH genes are associated with the accumulation of medium chain acylsugars, supporting our hypothesis above.

To further support our hypothesis, we extended our phenotypic analysis to six additional Solanaceae genera (fig. S8). According to our hypothesis, the presence of ACS and ECH genes should correlate with accumulation of medium chain acylsugars. Based on our phylogenetic and synteny analysis, the cluster with both *ACS* and *ECH* should only be present in the lineage after its divergence from the *Petunia* lineage. Consistent with our hypothesis, species such as *Jaltomata, Physalis, Iochroma, Atropa, and Hyoscyamus*, which are more closely related to *Solanum* species than they are to *Petunia*, have medium chain acylsugars (fig. S8). In contrast, *Salpiglossis sinuata*, diverged before the lineage leading to *Petunia* and *Solanum*, produces no detectable medium chain acylsugars (*28*) (fig. S8).

## Discussion

This study identified a *S. lycopersicum* chromosome 7 cluster of ACS, ECH, and BAHD acyltransferase genes including two involved in medium chain acylsugar biosynthesis. The discovery of this locus was prompted by our observation of increased C10 containing acylsugars in tomato recombinant lines carrying this region from the wild tomato *S. pennellii* chromosome 7. *In vitro* biochemistry revealed that Sl-AACS1 produces acyl-CoAs using C6-C12 fatty acids as substrates. The function of *AACS1* and *AECH1* in cultivated and wild tomato medium chain acylsugar biosynthesis was confirmed by genome editing, and extended to the phylogenetically distant *S. quitoense* using VIGS. The trichome tip cell-specific expression of these genes is similar to that of previously characterized acylsugar pathway genes (*24*).

There are increasing examples of plant specialized metabolic innovation evolving from gene duplication and neofunctionalization of primary metabolic enzymes (*1, 4, 47*). Recruitment of *Sl-AACS1* and *Sl-AECH1* from fatty acid metabolism provides new examples of ‘hijacking’ primary metabolic genes into acylsugar biosynthesis, in addition to an isopropylmalate synthase (*Sl-IPMS3*) and an invertase (*Sp-ASFF1*) (*32, 33*). We hypothesize that both AACS1 and AECH1 participate in generation of medium chain acyl-CoAs, the acyl donor substrates for ASAT-catalyzed acylsugar biosynthesis. Indeed, Sl-AACS1 exhibits *in vitro* function consistent with this hypothesis, efficiently utilizing medium chain fatty acids as substrates to synthesize acyl-CoAs. Strikingly, Sl-AECH1 perturbs membrane lipid composition when transiently expressed in *N. benthamiana* leaves, generating unusual C12-chain membrane lipids.

These results suggest that evolution of trichome tip cell-specific gene expression potentiated the co-option of *AACS1* and *AECH1* in medium chain acylsugar biosynthesis. Analogous to trichomes producing medium chain acylsugars, seeds of phylogenetically diverse plants accumulate medium chain fatty acid storage lipids (*48*). In contrast, fatty acids with unusual structures, including those of medium chain lengths, are rarely found in membrane lipids, presumably because these would perturb membrane bilayer structure and function (*49*). For example, seed embryo-specific expression of three neofunctionalized enzyme variant genes in *Cuphea* species – an acyl-ACP thioesterase (*50*), a 3-ketoacyl-ACP synthase (*51*), and a diacylglycerol acyltransferase (*52*) – lead to production of medium chain seed storage lipids (*53*). Trichome tip cell restricted expression of *AACS1* and *AECH1* represents an analogous example of diverting neofunctionalized fatty acid enzymes from general metabolism into cell-specific specialized metabolism. It is notable that we obtained evidence that Sl-AACS1 and Sl-AECH1 are targeted to mitochondria. Because the other characterized acylsugar biosynthetic enzymes – ASATs and Sl-IPMS3 – appear to be cytoplasmic, these results suggest that medium chain acylsugar substrates are intracellularly transported within the trichome tip cell.

Beyond employing functional approaches, this study demonstrates the value of a combined comparative genomic and metabolomic analysis in reconstructing the evolutionary history of a gene cluster: in this case over tens of millions of years. We propose that the acylsugar gene cluster started with a ‘founder’ BAHD acyltransferase gene, followed by sequential insertion of ECH and ACS (Fig. 6B). This *de novo* assembly process is analogous to evolution of the antimicrobial triterpenoid avenacin cluster in oat (*14, 54*). There are two noteworthy features of our approach. First, reconstructing acylsugar gene cluster evolution in a phylogenetic context allows us to deduce cluster composition in extant species (Fig. 6B). Second, it links cluster genotype with acylsugar phenotype and promotes prediction of acylsugar diversity across the Solanaceae (Fig. 6 and fig. S8): absence of either *ECH* or *ACS* in the synteny correlates with deficiency of medium chain acylsugars in the plant. The short chain acylsugar producer *Salpiglossis*, which diverged prior to the common ancestor of *Petunia*, validates the predictive power of this model.

The current architecture of the acylsugar cluster merely represents a snapshot of a dynamic genomic region. *De novo* assembly of the gene cluster was accompanied by gene duplication, transposition, pseudogenization, and deletion in different genera. In the extreme case of *Capsicum*, the cluster genes were deleted entirely (Fig. 6), which is consistent with lack of detectable trichome acylsugars in *Capsicum* plants. In *Nicotiana*, the ACH genes became pseudogenized (Fig. 6B) or poorly expressed (table S2), which is associated with lack of detectable plant medium chain acylsugars (fig. S8). In tomatoes, the trichome expressed *Solyc07g043660* is a tandem duplicate of *AACS1* (Fig. 2D), yet its deletion has no effect on trichome acylsugars (fig. S1D). A parsimonious explanation is that *Solyc07g043660* are experiencing functional divergence, which may eventually lead to pseudogenization as observed for other genes in the syntenic region.

The acylsugar gene cluster on chromosome 7 is syntenic to a chromosome 12 region containing *Sl-ASAT1* (fig. S2). This resembles the tomato steroidal alkaloid gene cluster consisting of eight genes that are dispersed into two syntenic chromosome regions (*17*). In fact, the tomato alkaloid cluster genes are directly adjacent to the acylsugar cluster, with six alkaloid genes on chromosome 7 and two on chromosome 12 (fig. S2). To our knowledge, this is the first discovery of two plant specialized metabolic gene clusters colocalizing in a genomic region, thus forming a ‘supercluster’. Tomato steroidal alkaloids and acylsugars both serve defensive roles in plants but are structurally distinct and stored in different tissues. This raises intriguing questions: how common are ‘superclusters’ encoding enzymes of distinct biosynthetic pathways in plant genomes; does this colocalization confer selective advantage through additive or synergistic effects of multiple classes of defensive metabolites? Answering these questions requires continued mining of metabolic gene clusters across broader plant species and analyzing the impacts of clustering in evolutionary and ecological contexts.

## Materials and methods

### Plant materials and trichome metabolite extraction

The seeds of cultivated tomato *Solanum lycopersicum* M82 were obtained from the C.M. Rick Tomato Genetic Resource Center (http://tgrc/ucdvis.edu). Tomato introgression lines (ILs) and tomato backcross inbred lines (BILs) were from Dr. Dani Zamir (Hebrew University of Jerusalem). The tomato seeds were treated with ½ strength bleach for 30 minutes and washed with de-ionized water three or more times before placing on wet filter paper in a Petri dish. After germination, the seedlings were transferred to peat-based propagation mix (Sun Gro Horticulture) and transferred to a growth chamber for two or three weeks under 16 h photoperiod, 28 °C day and 22 °C night temperatures, 50% relative humidity, and 300 μmol m^-2^ s^-1^ photosynthetic photon flux density. The youngest fully developed leaf was submerged in 1 mL extraction solution in a 1.5 mL screw cap tube and agitated gently for 2 min. The extraction solution contains acetonitrile/isopropanol/water (3:3:2) with 0.1% formic acid and 10 μM propyl-4-hydroxybenzoate as internal standard. The interactive protocol of acylsugar extraction is available in Protocols.io at http://dx.doi.org/10.17504/protocols.io.xj2fkqe.

### DNA construct assembly and tomato transformation

Assembly of the CRISPR-Cas9 constructs was as described (*32, 40*). Two guide RNAs (gRNAs) were designed targeting one or two exons of each gene to be knocked out by the CRISPR-Cas9 system. The gRNAs were synthesized gene (gBlocks) by IDT (Integrated DNA Technologies, location) (table S3). For each CRISPR construct, two gBlocks and four plasmids from Addgene, pICH47742::2x35S-5′UTR-hCas9 (STOP)-NOST (Addgene no. 49771), pICH41780 (Addgene no. 48019), pAGM4723 (Addgene no. 48015), pICSL11024 (Addgene no. 51144), were mixed for DNA assembly using the Golden Gate assembly kit (NEB).

For *in plant*a tissue specific reporter gene analysis, 1.8 kb upstream of the annotated translational start site of *Sl-AACS1* and *Sl-AECH* were amplified using the primer pairs SlAACS1-pro_F/R and SlECH1-pro_F/R (table S3). The amplicon was inserted into the entry vector pENTR/D-TOPO, followed by cloning into the GATEWAY vector pKGWFS7. For ectopically expressing *Sp-AACS1* in the cultivated tomato CRISPR mutant *sl-aacs1*, *Sp-AACS1* gene including 1.8 kb upstream of the translational start site of *Sp-AACS1* was amplified using the primer pair SpAACS1-pro-gene_F/R (table S3). The amplicon was inserted into the entry vector pENTR/D-TOPO, followed by cloning into the GATEWAY vector pK7WG.

The plant transformation of *S. lycopersicum* M82 and *S. pennellii* LA0716 was performed using the *Agrobacterium tumefaciens* strain AGL0 following published protocols (*32, 55*). The primers used for genotyping the *S. lycopersicum* M82 transgenic plants harboring pK7WG or pKGWFS7 construct are listed in table S3. For genotyping the *S. lycopersicum* M82 CRISPR mutants in the T1 generation, the sequencing primers listed in table S3 were used to amplify the genomic regions harboring the gRNAs and the resultant PCR products were sent for Sanger sequencing. For genotyping the *S. pennellii* LA0716 CRISPR mutants in the T0 generation, the sequencing primers listed in table S3 were used to amplify the genomic regions harboring the gRNAs. The resulting PCR products were cloned into the pGEM-T easy vector (Promega) and transformed into *E. coli*. Plasmids from at least six individual *E. coli* colonies containing the amplified products were extracted and verified by Sanger sequencing.

### Protein subcellular targeting in tobacco mesophyll cells

For protein subcellular targeting analysis, the open reading frame (ORF) of *Sl-AACS1*, *Sl-AECH*, and *Solyc07g043660* were amplified using the primers listed in table S3. These amplicons were inserted into pENTR/D-TOPO respectively, followed by subcloning into the GATEWAY vectors pEarleyGate102 (no. CD3-684) and pEarleyGate104 (no. CD3-686), which were obtained from Arabidopsis Biological Resource Center (ABRC). For the pEarleyGate102 constructs, the CFP was fused to the C-terminal of the tested proteins. For the pEarleyGate104 constructs, the YFP was fused to the protein N-terminus. Transient expressing the tested proteins was performed following an established protocol (*56*) with minor modifications. In brief, cultures of *A. tumefaciens* (strain GV3101) harboring the expression vectors were washed and resuspended with the infiltration buffer (20 mM acetosyringone, 50 mM MES pH 5.7, 0.5% glucose [w/v] and 2 mM Na_3_PO_4_) to reach OD_600nm_ = 0.05. Four-week-old tobacco (*Nicotiana tabacum* cv. Petit Havana) plants grew in 21⃞ and 8 h short-day conditions were infiltrated, and then maintained in the same growth condition for three days before being sampled for imaging. The GV3101 cultures containing the mitochondria marker MT-RFP (*42*) were co-infiltrated to provide the control signals for mitochondrial targeting. In separate experiments, the GV3101 cultures containing the peroxisome marker RFP-PTS (*42*) were co-infiltrated to provide the control signals for peroxisomal targeting.

### Confocal Microscopy

A Nikon A1Rsi laser scanning confocal microscope and Nikon NIS-Elements Advanced Research software were used for image acquisition and handling. For visualizing GFP fluorescence in trichomes of the tomato transformants, the excitation wavelength at 488 nm and a 505- to 525-nm emission filter were used for the acquisition. For visualizing signals of fluorescence proteins in the tobacco mesophyll cells, CFP, YFP and RFP, respectively, were detected by excitation lasers of 443 nm, 513 nm, 561 nm and emission filters of 467-502 nm, 521-554 nm, 580-630 nm.

### *N. benthamiana* transient gene expression and membrane lipid analysis

For *N. benthamiana* transient expression of *Sl-AACS1*, *Sl-AECH1*, and homologs of *AECH1*, the ORFs of these genes were amplified using primers listed in table S3, followed by subcloning into pEAQ-HT vector using the Gibson assembly kit (NEB). Linearization of pEAQ-HT vector was performed by XhoI and AgeI restriction enzyme double digestion. *A. tumefaciens* (strain GV3101) harboring the pEAQ-HT constructs were grown in LB medium containing 50 µg/mL kanamycin, 30 µg/mL gentamicin, and 50 µg/ml rifampicin at 30 ⃞. The cells were collected by centrifugation at 5000 g for 5 min and washed once with the resuspension buffer (10 mM MES buffer pH 5.6, 10 mM MgCl_2_, 150 µM acetosyringone). The cell pellet was resuspended in the resuspension buffer to reach OD_600nm_ = 0.5 for each strain and was incubated at room temperature for 1 h prior to infiltration. Leaves of 4 to 5-week-old *N. benthamiana* grown under 16 h photoperiod were used for infiltration. Five days post infiltration, the infiltrated leaves were harvested, ground in liquid nitrogen, and stored at -80 ⃞ for later analysis.

The membrane lipid analysis was performed as previously described (*57*). In brief, the *N. benthamiana* leaf polar lipids were extracted in the organic solvent containing methanol, chloroform, and formic acid (20:10:1, v/v/v), separated by thin layer chromatography (TLC), converted to fatty acyl methylesters (FAMEs), and analyzed by gas-liquid chromatography (GLC) coupled with flame ionization. The TLC plates (TLC Silica gel 60, EMD Chemical) were activated by ammonium sulfate before being used for lipid separation. Iodine vapor was applied to TLC plates after lipid separation for brief reversible staining. Different lipid groups on the TLC plates were marked with a pencil and were scraped for analysis. For LC/MS analysis, lipids were extracted using the buffer containing acetonitrile/isopropanol/water (3:3:2) with 0.1% formic acid and 10 μM propyl-4-hydroxybenzoate as the internal standard.

### Protein expression and ACS enzyme assay

To express His-tagged recombinant protein Sl-AACS1, the full-length *Sl-AACS* ORF sequence was amplified using the primer pair SlAACS1-pET28_F/R (table S3) and was cloned into pET28b (EMD Millipore) using the Gibson assembly kit (NEB). The pET28b vector was linearized by digesting with BamHI and XhoI to create overhangs compatible for Gibson assembly. The pET28b constructs were transformed into BL21 Rosetta cells (EMD Millipore). The protein expression was induced by adding 0.05 mM isopropyl *β*-D-1-thiogalactopyranoside to the cultures when the OD_600nm_ = 0.5. The *E. coli* cultures were further grown overnight at 16 ⃞ 120 rpm. The His-tagged proteins were purified by Ni-affinity gravity-flow chromatography using the Ni-NTA agarose (Qiagen) following the product manual.

Measurement of acyl-CoA synthetase activity was performed using minor modifications of the coupled enzyme assay described by Schneider et al. (*58*). A multimode plate reader (PerkinElmer, mode EnVision 2104) compatible with the 96-well UV microplate was used for the assays. The fatty acid substrates were dissolved in 5% Triton X-100 (v/v) to make 5 mM stock solutions. The enzyme assay premix was prepared containing 0.1 M Tris-HCl (pH 7.5), 2 mM dithiothreitol, 5 mM ATP, 10 mM MgCl_2_, 0.5 mM CoA, 0.8 mM NADH, 250 µM fatty acid substrate, 1 mM phosphoenolpyruvate, 20 units myokinase (Sigma-Aldrich, catalog no. M3003), 10 units pyruvate kinase (Roche, 10128155001), 10 units lactate dehydrogenase (Roche, 10127230001), and was aliquoted 95 µL each to the 96-well microplate. The reaction was started by adding 5 µL (1-2 µg) proteins. The chamber of the plate reader was set to 30 ⃞ and the OD at 340 nm was recorded every 5 min for 40 min. Oxidation of NADH, which is monitored by the decrease of OD_340nm_, was calculated using the NADH extinction coefficient 6.22 cm^2^ µmol^-1^. Every two moles of oxidized NADH is equivalent to one mole of acyl-CoA product generated in the reaction. To measure the parameters of enzyme kinetics, the fatty acid substrate concentration was varied from 0 to 500 µM, with NADH set at 1 mM. The fatty acid substrates, sodium acetate (C2:0), sodium butyrate (C4:0), sodium hexanoate (C6:0), sodium octanoate (C8:0), sodium decanoate (C10:0), sodium laurate (C12:0), sodium myristate (C14:0), sodium palmitate (C16:0), and sodium stearate (C18:0), were purchased from Sigma-Aldrich. *Trans*-2-decenoic acid (C10:1), 8-methylnonanoic acid (iC10:0), 3-hydroxy lauric acid (C12:OH), and 3-hydroxy myristic acid (C14:OH) were purchased from Cayman Chemical.

### RNA extraction, sequencing, and differential gene expression analysis

Total RNA was extracted from trichomes isolated from stems and shaved stems of 7-week-old *S. pennellii* LA0716 plants using the RNAeasy Plant Mini kit (Qiagen) and digested with DNase I. A total of four RNA samples extracted from two tissues with two replicates were used for RNA sequencing. The sequencing libraries were prepared using the KAPA Stranded RNA-Seq Library Preparation Kit. Libraries went through quality control and quantification using a combination of Qubit dsDNA high sensitivity (HS), Applied Analytical Fragment Analyzer HS DNA and Kapa Illumina Library Quantification qPCR assays. The libraries were pooled and loaded onto one lane of an Illumina HiSeq 4000 flow cell. Sequencing was done in a 2x150bp paired end format using HiSeq 4000 SBS reagents. Base calling was done by Illumina Real Time Analysis (RTA) v2.7.6 and output of RTA was demultiplexed and converted to FastQ format with Illumina Bcl2fastq v2.19.1.

The paired end reads were filtered and trimmed using Trimmomatic v0.32 (*59*) with the setting (LLUMINACLIP: TruSeq3-PE.fa:2:30:10 LEADING:3 TRAILING:3 SLIDINGWINDOW:4:30), and then mapped to the *S. pennellii* LA0716 genome v2.0 (*60*) using TopHat v1.4 (*61*) with the following parameters: -p (threads) 8, -i (minimum intron length) 50, -I (maximum intron length) 5000, and -g (maximum hits) 20. The FPKM (Fragments Per Kilobase of transcript per Million mapped reads) values for the genes were analyzed via Cufflinks v2.2 (*62*).For differential expression analysis, the HTseq package (*63*) in Python was used to get raw read counts, then Edge R version 3.22.5 (*64*) was used to compare read counts between trichome-only RNA and shaved stem RNA using a generalized linear model (glmQLFit).

### VIGS and qRT-PCR

For VIGS analysis of *Sq-AACS1* and *Sq-AECH1* in *S. quitoense*, the fragments of these two genes, as well as the phytoene desaturase (PDS) gene fragment, were amplified using the primers listed in table S3, cloned into pTRV2-LIC (ABRC no. CD3-1044) using the ligation-independent cloning method (*65*), and transformed into *A. tumefaciens* (strain GV3101). The VIGS experiments were performed as described (*66*). In brief, the Agrobacterium strains harboring pTRV2 constructs, the empty pTRV2, or pTRV1 were grown overnight in separate LB cultures containing 50 µg/mL kanamycin, 10 µg/mL gentamicin, and 50 µg/ml rifampicin at 30 ⃞. The cultures were re-inoculated in the induction media (50 mM MES pH5.6, 0.5% glucose [w/v], 2 mM NaH_2_PO_4_, 200 µM acetosyringone) for overnight growth. The cells were harvested, washed, and resuspended in the buffer containing 10 mM MES, pH 5.6, 10 mM MgCl_2_, and 200 µM acetosyringone with the OD_600nm_ = 1. Different cultures containing pTRV2 constructs were mixed with equal volume of pTRV1 cultures prior to infiltration. The 2- to 3-week-old young *S. quitoense* seedlings grown under 16 h photoperiod at 24 °C were used for infiltration: the two fully expanded cotyledons were infiltrated. Approximately three weeks post inoculation, the fourth true leaf of each infiltrated plant was cut in half for acylsugar quantification and gene expression analysis, respectively. The onset of the albino phenotype of the control group infiltrated with the PDS construct was used as a visual marker to determine the harvest time and leaf selection for the experimental groups. At least fourteen plants were analyzed for each construct. The trichome acylsugars were extracted using the solution containing acetonitrile/isopropanol/water (3:3:2) with 0.1% formic acid and 1 μM telmisartan as internal standard, following the protocol at http://dx.doi.org/10.17504/protocols.io.xj2fkqe.

The leaf RNA was extracted using RNeasy Plant Mini kits (Qiagen) and digested with DNase I. The first-strand cDNA was synthesized by Superscript II (Thermofisher Scientific) using total RNA as templates. Quantitative real-time PCR was performed to analyze the *Sq-AACS1* or *Sq-AECH1* mRNA in *S. quitoense* leaves using the primers listed in table S3. EF1α was used as a control gene. A Quantstudio 7 Flex Real-Time PCR System with Fast SYBR Green Master Mix (Applied Biosystems) was used for the analysis. The relative quantification method (2^-ΔΔct^) was used to evaluate the relative transcripts levels.

### LC/MS analysis

Trichome acylsugars extracted from tomato IL and BILs were analyzed using a Shimadzu LC-20AD HPLC system connected to a Waters LCT Premier ToF mass spectrometer. Ten microliter samples were injected into a fused core Ascentis Express C18 column (2.1 mm × 10 cm, 2.7 μm particle size; Sigma-Aldrich) for reverse-phase separation with column temperature set at 40 °C. The starting condition was 90% solvent A (0.15% formic acid in water) and 10% solvent B (acetonitrile) with flow rate set to 0.4 mL/min. A 7 min linear elution gradient was used: ramp to 40% B at 1 min, then to 100% B at 5 min, hold at 100% B to 6 min, return to 90% A at 6.01 min and hold until 7 min.

For analyzing trichome acylsugars extracted from *S. pennellii* transgenic plants and membrane lipids from *N. benthamiana*, a Shimadzu LC-20AD HPLC system connected to a Waters Xevo G2-XS QToF mass spectrometer was used. The starting condition were 95% solvent A (10 mM ammonium formate, pH 2.8) and 5% solvent B (acetonitrile) with flow rate set to 0.3 mL/min. A 7 min linear elution gradient used for acylsugar analysis was: ramp to 40% B at 1 min, then to 100% B at 5 min, hold at 100% B to 6 min, return to 95% A at 6.01 min and hold until 7 min. A 12 min linear elution gradient used for the lipid analysis was: ramp to 40% B at 1 min, then to 100% B at 5 min, hold at 100% B to 11 min, return to 95% A at 11.01 min and hold until 12 min.

For analyzing trichome acylsugars extracted from other plants, a Waters Acquity UPLC was coupled to a Waters Xevo G2-XS QToF mass spectrometer. The starting condition was 95% solvent A (10 mM ammonium formate, pH 2.8) and 5% solvent B (acetonitrile) with flow rate set to 0.3 mL/min. A 7 min linear elution gradient was: ramp to 40% B at 1 min, then to 100% B at 5 min, hold at 100% B to 6 min, return to 95% A at 6.01 min and held until 7 min. A 14 min linear elution gradient was: ramp to 35% B at 1 min, then to 85% B at 12 min, then to 100% B at 12.01 min, hold at 100% B to 13 min, return to 95% A at 13.01 min and held until 14 min.

For Waters LCT Premier ToF mass spectrometer, the MS settings of electrospray ionization in negative mode were: 2.5 kV capillary voltage, 100°C source temperature, 350°C desolvation temperature, 350 liters/h desolvation nitrogen gas flow rate, 10 V cone voltage, and mass range *m/z* 50 to 1500 with spectra accumulated at 0.1 seconds/function. Three collision energies (10, 40, and 80 eV) were used in separate acquisition functions to generate both molecular ion adducts and fragments. For Waters Xevo G2-XS QToF mass spectrometer, the MS settings of the negative ion-mode electrospray ionization were as follows: 2.00 kV capillary voltage, 100 °C source temperature, 350 °C desolvation temperature, 600 liters/h desolvation nitrogen gas flow rate, 35V cone voltage, mass range of *m/z* 50 to 1000 with spectra accumulated at 0.1 seconds/function. Three collision energies (0, 15, and 35 eV) were used in separate acquisition functions. The MS settings for positive ion-mode electrospray ionization were: 3.00 kV capillary voltage, 100 °C source temperature, 350 °C desolvation temperature, 600 liters/h desolvation nitrogen gas flow rate, 35V cone voltage, mass range of *m/z* 50 to 1000 with spectra accumulated at 0.1 seconds/function. Three collision energies (0, 15, and 45 eV) were used in separate acquisition functions. The Waters Quanlynx software was used to integrate peak areas of the selected ion relative to the internal standard. For quantification purpose, data collected with the lowest collision energy was used in the analysis.

### Synteny scan

Protein sequences of annotated genes and the corresponding annotation files in General Feature Format (GFF) of 11 Solanaceae species, *Ipomoea trifida,* and *Coffea canephora* were downloaded from National Center for Biotechnology Information (NCBI, https://www.ncbi.nlm.nih.gov/genome/) or Solanaceae Genomics Network (SGN, https://solgenomics.net/). The GFF files contain the coordinates of annotated genes on assembled chromosomes or scaffolds. The sources and version numbers of sequences and GFF files used are: *S. lycopersicum* ITAG3.2 (SGN) and V2.5 (NCBI), *S. pennellii* SPENNV200 (NCBI) and v2.0 (SGN), *S. tuberosum* V3.4 (SGN), *S. melongena* r2.5.1 (SGN), *Capsicum annuum L. zunla-1* V2.0 (SGN), *C. annuum_var. glabriusculum* V2.0 (SGN), *Nicotiana attenuata* NIATTr2 (SGN), *N. tomentosiformis* V01 (NCBI), *N. benthamiana* V1.0.1 (SGN), *Petunia axillaris* V1.6.2 (SGN), *P. inflata* V1.0.1 (SGN), *I. trifida* V1.0 (NCBI), and *C. canephora* Vx (SGN).

To hypothesize the evolutionary history of genes in the acylsugar gene cluster, putative pseudogenes, which are homologs to protein-coding genes but with predicted premature stops/frameshifts and/or protein sequence truncation, were also identified for each species as described (*67*). Genome-wide syntenic analysis was conducted using annotated protein-coding genes and putative pseudogenes from all the species with MCScanX-transposed (*68*) as described (*67*). The MCScanX-based analysis did not lead to a syntenic block of acylsugar gene cluster on chromosome 7 of *S. melongena* r2.5.1, which can be due to true absence of the gene cluster, issues with genome assembly, or lack of coverage. To verify this, protein sequences of *S. lycopersi*cum genes in genomic blocks on chromosome 7 were searched against an updated *S. melongena* genome from The Eggplant Genome Project (http://ddlab.dbt.univr.it/eggplant/) that led to the identification of the gene cluster.

### Acylsugar acyl chain composition analysis by GC-MS

Acyl chains were characterized from the corresponding fatty acid ethyl esters following transesterification of acylsugar extractions as previously reported (*33*). Plants were grown for 4-8 weeks and approximately ten leaves were extracted for 3 minutes in 10 mL of 1:1 isopropanol:acetonitrile with 0.01% formic acid. Extractions were dried to completeness using a rotary evaporator and then 300 µL of 21% (v/v) sodium ethoxide in ethanol (Sigma) was added and incubated for 30 minutes with gentle rocking and vortexing every five minutes and 400 µL hexane was added and vortexed for 30 seconds. To the hexane layer, 500 µL of saturated sodium chloride in water was added and vortexed to facilitate a phase separation. After phase separation, the top hexane layer was transferred to a new tube. The phase separation by addition of 500 µL hexane was repeated twice, with the final hexane layer transferred to a 2 mL glass vial with a glass insert.

The fatty acid ethyl esters were analyzed using an Agilent 5975 single quadrupole GC-MS equipped with a 30-m, 0.25-mm internal diameter fused silica column with a 0.25-µm film thickness VF5 stationary phase (Agilent). Injection of 1 µL of each hexane extract was performed using splitless mode. The gas chromatography program was as follows: inlet temperature, 250°C; initial column temperature, 70°C held for 2 min; ramped at 20°C/min until 270°C, then held at 270°C 3 min. The helium carrier gas was used with 70 eV electron ionization. Acyl chain type was determined through NIST Version 2.3 library matches of the mass spectra of the corresponding ethyl ester and relative abundances were determined through integrating the corresponding peak area over the total acyl chain peak area.

## Acknowledgements

We thank the C.M. Rick Tomato Genetics Resource Center (University of California Davis, CA USA) for providing tomato seeds, Zamir lab in Hebrew University of Jerusalem for providing tomato ILs and BILs seeds. We acknowledge Dr. Kun Wang and Dr. Christoph Benning for their helpful guidance in lipid analysis. We thank Krystle Wiegert-Rininger and Cornelius Barry for their help in RNA sequencing and Dr. Kent Chapman from University of North Texas for helpful discussions. We acknowledge Kathleen Imre and Sara Haller for their help with tomato transformation. We thank the MSU Center for Advanced Microscopy and RTSF Mass Spectrometry and Metabolomics Core Facilities for their support with LC/MS analysis.

## Author contribution

P.F. and R.L.L. conceived the study. P.F., P.W., Y.R.L, B.J.L., B.M.M, C.A.S., R.C., P.C., F.B., S.H.S, and R.L.L. designed the experiments and analyzed the data. P.F., Y.R.L, B.J.L., C.A.S., R.C., and P.C. performed the experiments. P.F. and R.L.L wrote the first draft. All authors contributed to the final version of the manuscript.

## Data and materials availability

All data needed to evaluate the conclusions in the paper are present in the paper and/or the Supplementary Materials. The RNA-seq reads were deposited in the National Center for Biotechnology Information Sequence Read Archive under the accession number PRJNA605501. Sequence data used in this study are in the GenBank/EMBL data libraries under these accession numbers: *Sl-AACS1*(MT078737), *Sl-AECH1*(MT078736), *Sp-AACS1*(MT078735), *Sp-AECH1*(MT078734), *Sq-AACS1*(MT078732), *Sq-AECH1*(MT078731), *Sq_c35719* (MT078733). The following materials require a material transfer agreement: pEAQ-HT, pK7WG, pKGWFS7, pEarleyGate102, pEarleyGate104, pTRV2-LIC, pICH47742::2x35S-5’UTR-hCas9(STOP)-NOST, pICH41780, pAGM4723, and pICSL11024. Requests for biological materials or data should be submitted to R.L.L. at.

## Supplementary Materials

**Fig. S1.**
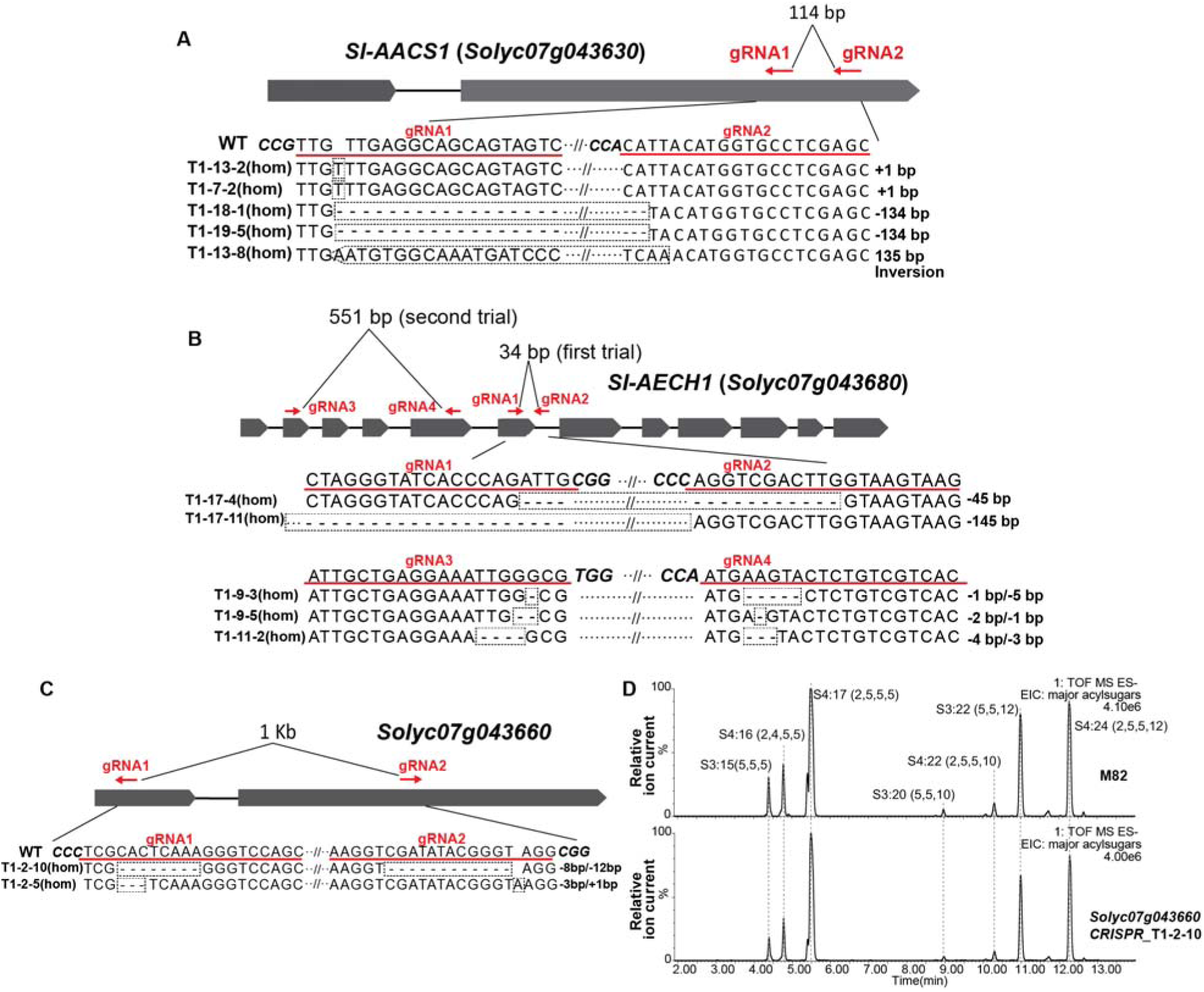
CRISPR-Cas9-mediated gene knockouts in cultivated tomato *S. lycopersicum*. The gRNAs targeting *Sl-AACS1* (**A**), *Sl-AECH1*(**B**), and *Solyc07g043660* (**C**) in cultivated tomato are highlighted with red lines and text. Two pairs of gRNAs were designed that target *Sl-AECH1*, which were followed by two trials of plant transformation. DNA sequences of the self-crossed T1 generation transgenic lines carrying homozygous gene edits are shown beneath the gene model. Dotted rectangle boxes highlight edited sequences. (**D**) Electrospray ionization negative (ESI^-^) mode, LC/MS extracted ion chromatograms of seven major trichome acylsugars of the CRISPR mutant *solyc07g043660* and the M82 parent.

**Fig. S2.**
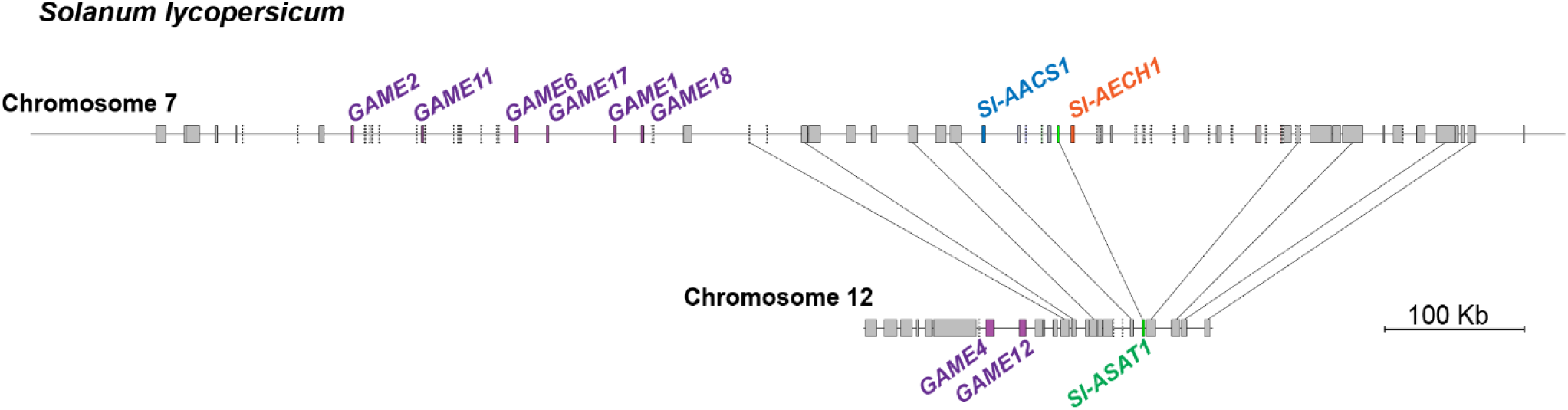
The ‘supercluster’ syntenic region of cultivated tomato *S. lycopersicum* chromosome 7 and 12 harboring the acylsugar and steroidal glycoalkaloid gene clusters. The gene models are represented by rectangles, with pseudogenes labeled with dotted lines. The lines linking the gene models of the two chromosomes denote putative orthologous genes in the synteny. GAME genes involved in glycoalkaloid metabolism were previously reported (*17*).

**Fig. S3.**
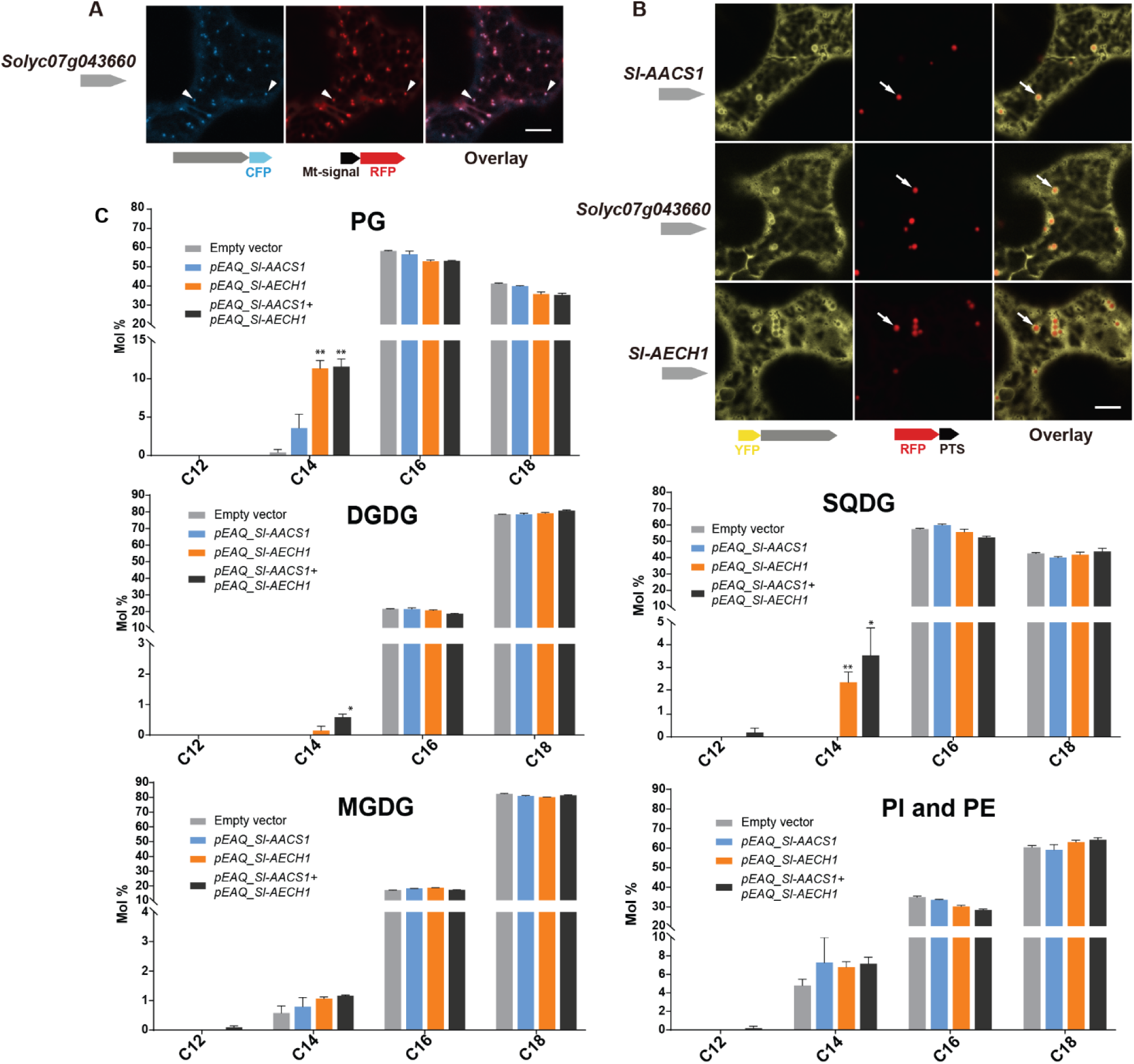
Characterization of cluster genes using leaf transient expression: protein subcellular targeting and impacts on lipid metabolism. (**A**) Confocal images of co-expression analysis in *N. tabacum* leaf epidermal cells using C-terminal CFP-tagged Solyc07g043660 and the mitochondrial marker MT-RFP. Arrowheads point to mitochondria that are indicated by MT-RFP fluorescent signals. White bar equals 10 μm. (**B**) Confocal images of co-expression analysis in *N. tabacum* leaf epidermal cells using N-terminal YFP-tagged Sl-AACS1, Sl-AECH1, or Solyc07g043660, and the peroxisomal marker RFP-PTS. Arrowheads point to peroxisomes that are indicated by RFP-PTS fluorescent signals. Bar equals 10 μm. (**C**) *N. benthamiana* leaf membrane lipid acyl chain composition. The results from infiltrating *Sl-AACS1* or *Sl-AECH1* individually, and infiltrating *Sl-AECH1* and *Sl-AACS1* together are shown. The lipid abbreviations are: phosphatidylglycerol (PG), sulfoquinovosyl diacylglycerol (SQDG), digalactosyldiacylglycerol (DGDG), monogalactosyldiacylglycerol (MGDG), phosphtatidylinositols (PI), and phosphtatidylethanolamine (PE). Mole percentage (Mol %) of the acyl chains from membrane lipids with carbon number 12, 14, 16, and 18 are shown for three biological replicates ± SE. *p < 0.05, **p < 0.01. Welch two-sample *t* test was performed comparing each experimental with the empty vector control. PI and PE lipids were adjacent on the TLC plates and were pooled for analysis. Acyl groups of the same chain lengths with saturated and unsaturated bonds were combined in the calculation.

**Fig. S4.**
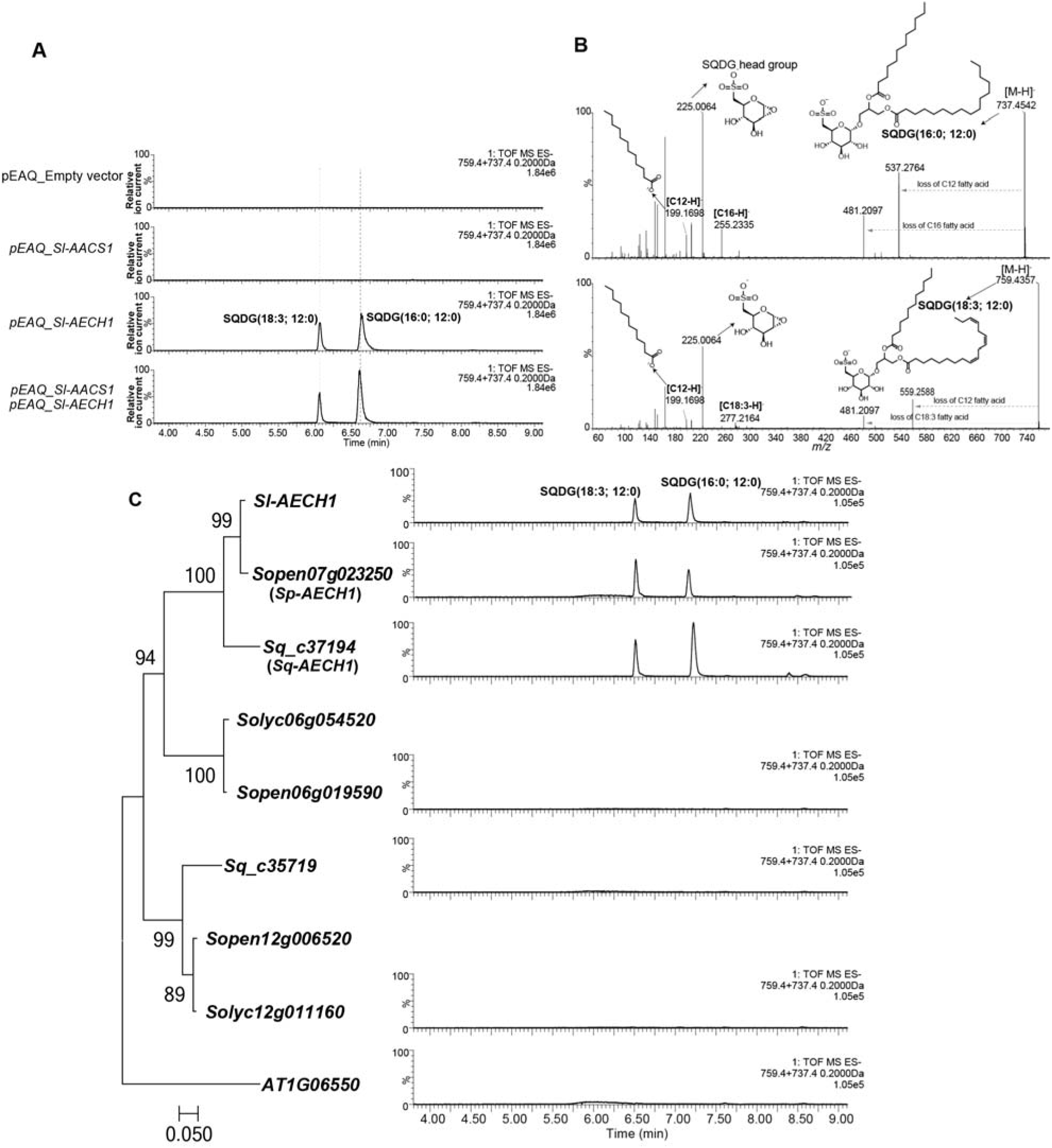
The closest homologs of *Sl-AECH1* from other *Solanum* species generate medium chain lipids when transiently expressed in *N. benthamiana*. (**A**) ESI^-^ mode, LC/MS extracted ion chromatograms of two SQDGs with C12 as peaks diagnostic of lipids containing medium chain fatty acids. (**B**) Mass spectra of two SQDGs contain C12 acyl chain. Fragmentation of SQDG (16:0, 12:0) and SQDG (18:3, 12:0) in ESI^-^ mode revealed the fragment ion C12 fatty acid (*m/z*: 199.17) and the SQDG head group (*m/z*: 225.0). (**C**) Among the putative *Sl-AECH1* homologs tested, *Sopen07g023250* (*Sp-AECH1*) and *Sq_c37194* (*Sq-AECH1*), from *S. pennellii* and *S. quitoense* respectively, generated medium chain SQDG in the infiltrated leaves. The maximum likelihood phylogenetic gene tree shown on the left was generated using the nucleotide sequences of ECH-annotated variants, with the Arabidopsis *AT1G06550* serving as an outgroup. The bootstrap values were obtained with 1000 replicates.

**Fig. S5.**
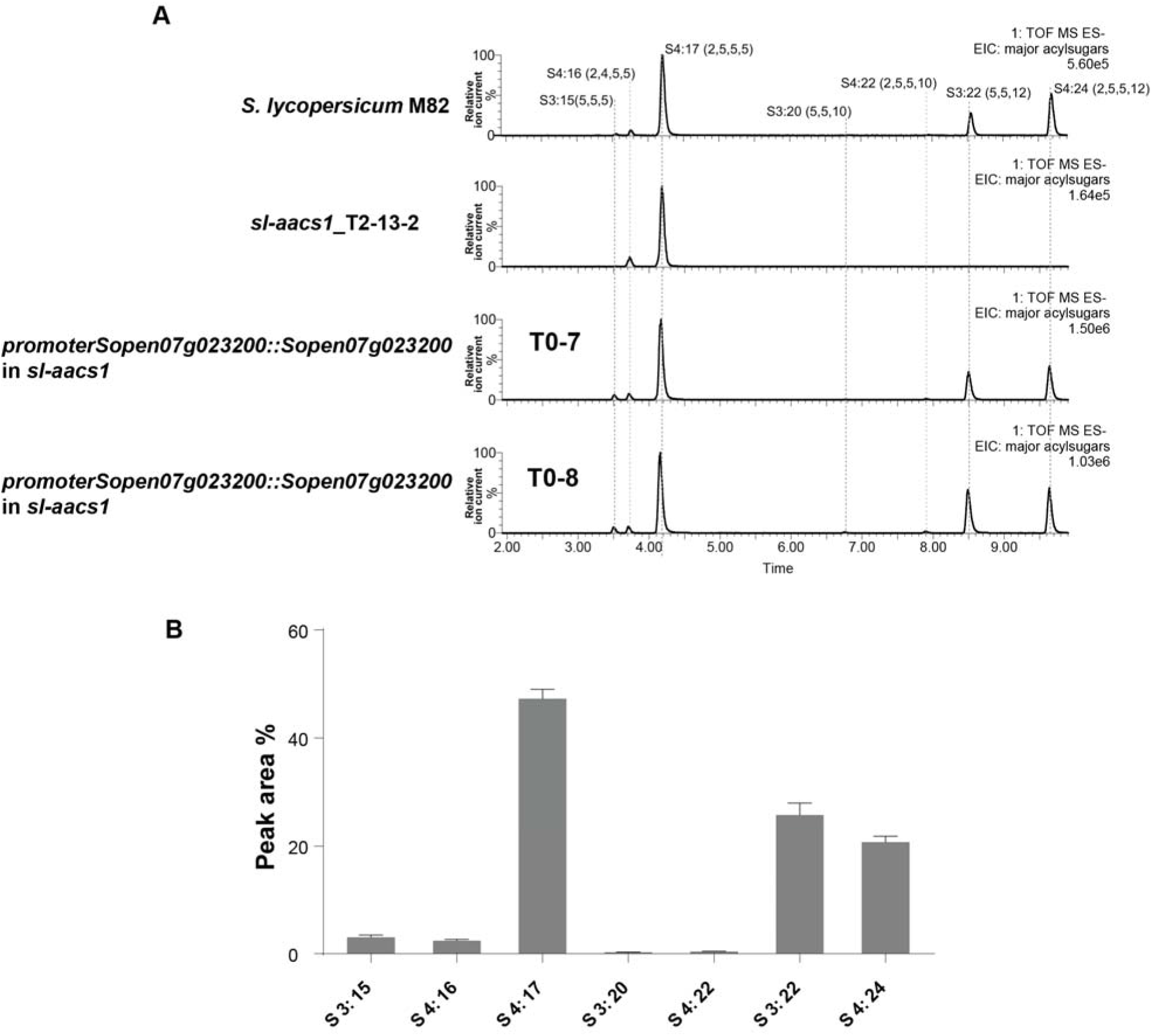
Stable *Sp-AACS1* transformation of the M82 CRISPR mutant *sl-aacs1* restores C12 containing acylsugars. (**A**) ESI^-^ mode, LC/MS extracted ion chromatograms are shown for seven major acylsugar peaks extracted from trichome of *S. lycopersicum* M82, *sl-aacs1*, and two independent T0 generation suppressed *sl-aacs1* transgenic lines expressing *Sp-AACS1* under its own promoter. (**B**) Peak area percentage of seven major trichome acylsugars of the suppression transgenic plants. The sum of the peak area percentage of each acylsugar equals to 100%. The results of acylsugar peak area percentage were calculated from six independent T0 transgenic lines ± SE.

**Fig. S6.**
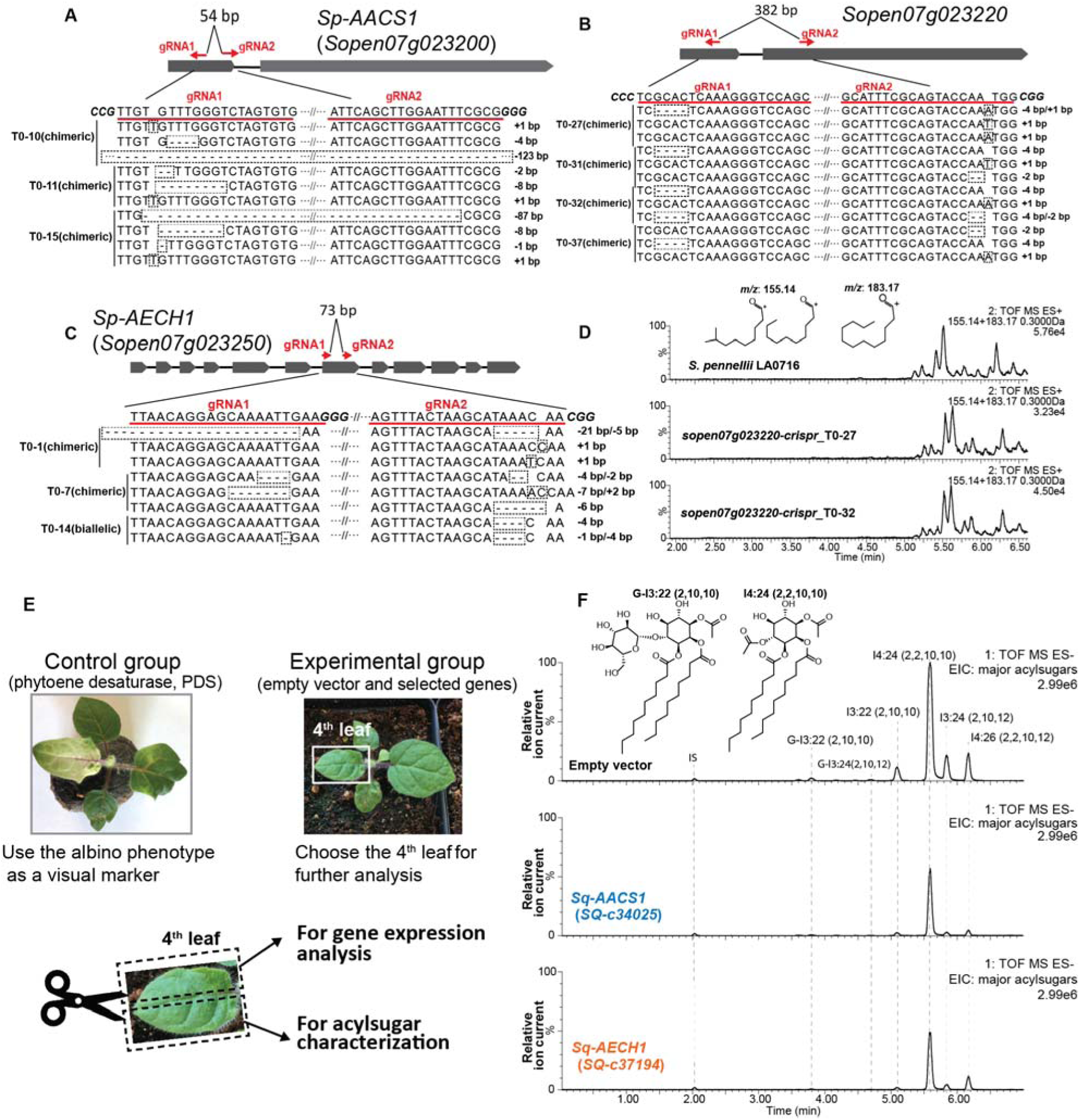
Functional analysis of *AACS1* and *AECH1* in *S. pennellii* and *S. quitoense* via CRISPR-Cas9 system and VIGS, respectively. The design of gRNAs targeting wild tomato *S. pennellii* LA0716 *Sp-AACS1* (**A**), *Sopen07g023250* (**B**), and *Sp-AECH1* (**C**) is highlighted with red lines and text. The transgenic T0 generation carrying chimeric or biallelic gene edits are shown beneath the gene model. DNA sequence of the gene edits was obtained through Sanger sequencing of cloned plant DNA fragments. The gene edits are highlighted with dotted rectangular boxes. (**D**) ESI^+^ mode, LC/MS extracted ion chromatograms shown for C10 (*m/z*: 155.14) and C12 (*m/z*: 183.17) fatty acid ions corresponding to medium chain trichome acylsugars extracted from the CRISPR mutants *sopen07g023220* and the *S. pennellii* LA0716 parent. (**E**) Experimental design of VIGS in *S. quitoense*. A control group silencing the PDS genes was performed in parallel with the experimental groups. The onset of the albino phenotype of the control group was used as a visual marker to determine the harvest time and leaf selection in the experimental groups. The fourth true leaves were harvested and cut in half for gene expression analysis and acylsugar quantification, respectively. (**F**) ESI^-^ mode, LC/MS extracted ion chromatograms of six major acylsugars of *S. quitoense* for the experimental group. The three LC/MS chromatograms show representative acylsugar profiles of the empty vector control plants and the VIGS plants targeting *Sq-AACS1* and *Sq-AECH1*.

**Fig. S7.**
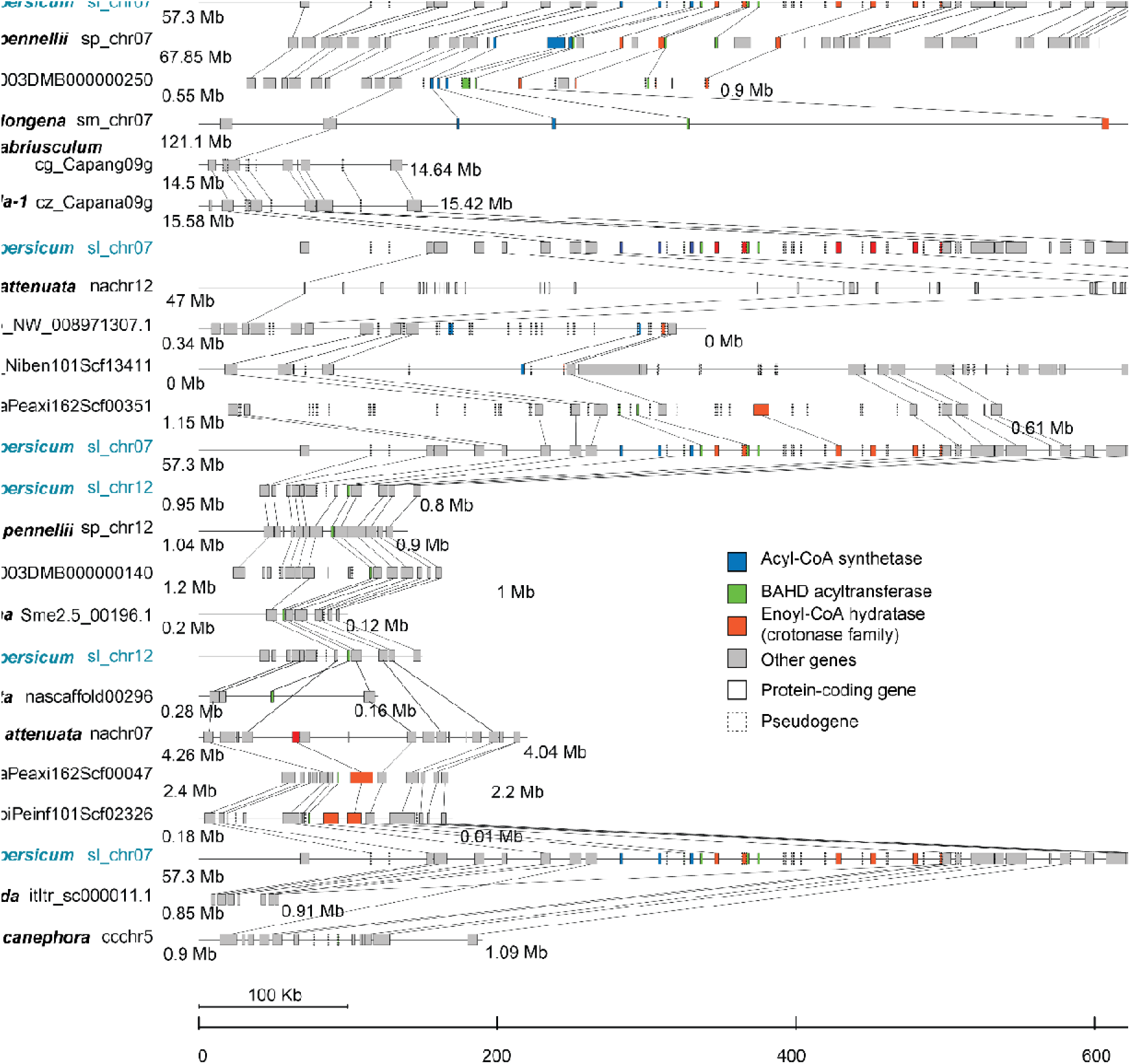
Syntenic regions containing the acylsugar gene cluster. The species name and chromosome/scaffold identifier are indicated with *S. lycopersinum* in blue font. Rectangle: protein-coding gene (solid line) or pseudogene (dotted line) colored according to the type of genes. Line connecting two genes: putative orthologous genes. Numbers underneath chromosomes: chromosome coordinates in million bases (Mb).

**Fig. S8.**
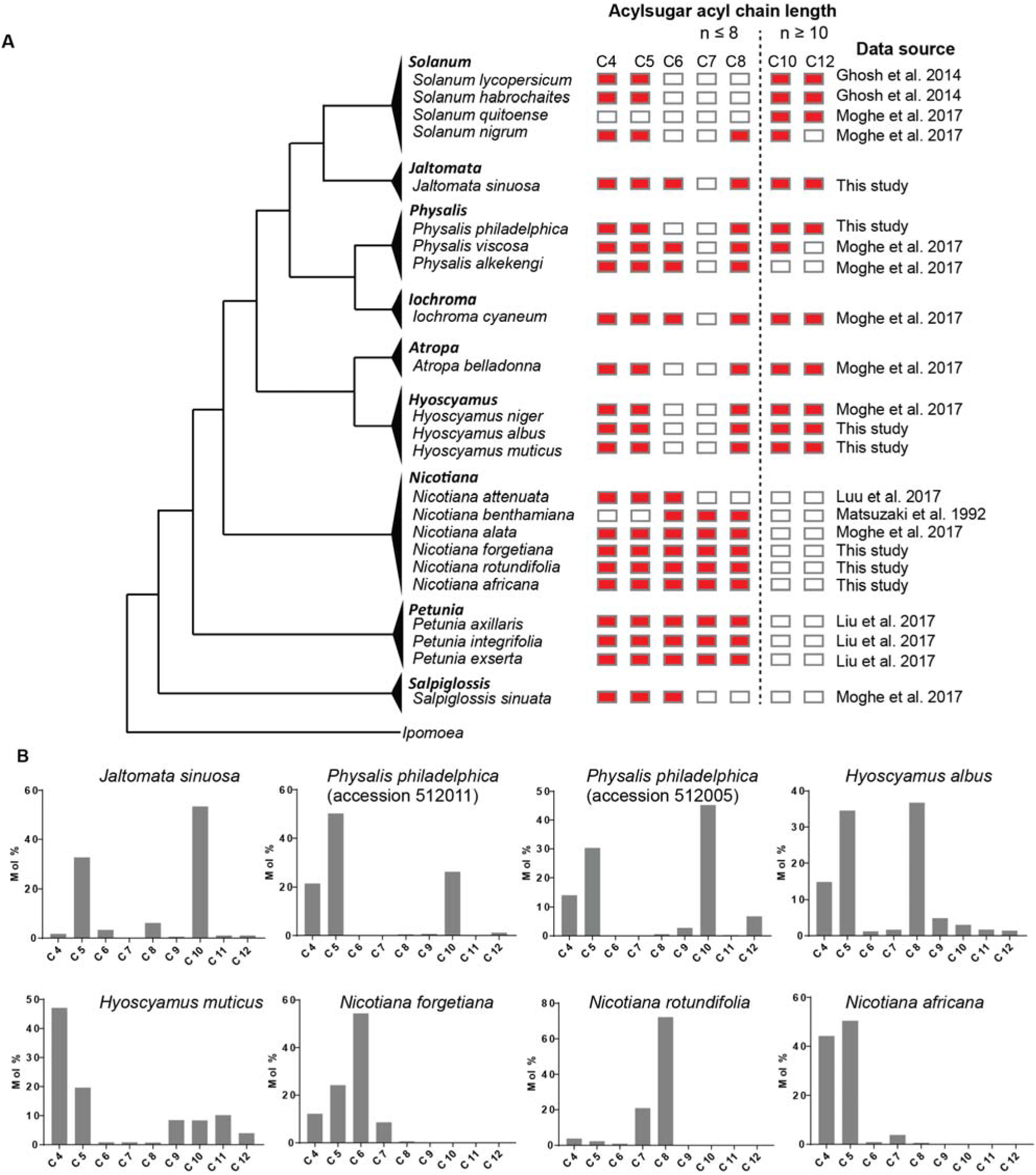
Phylogenetic distribution of acylsugar acyl chains with different lengths across the Solanaceae family. (**A**) The collated results of acylsugar acyl chain distribution across different Solanaceae species. Red rectangles indicate detectable acyl chains in the acylsugars produced in the tested species and white rectangles indicate no detectable signals. The data source where the results are derived is listed on the right. (**B**) Results of acylsugar acyl chain characterization of selected Solanaceae species. Mole percentage (Mol %) of acylsugar acyl chains with different lengths were obtained from GC/MS analysis of fatty acid ethyl esters.

### Other Supplementary Materials not in this file

**Table S1.** Gene expression levels of all analyzed transcripts in *S. pennellii* LA0716.

**Table S2.** The relative transcript abundance of ACS- and ECH-annotated genes in the syntenic region of *Nicotiana attenuate*.

**Table S3.** Synthesized gene fragments and primers used in this study.

